# Serum amyloid A3 deficiency modulates aortic immune composition and attenuates murine atherosclerosis progression

**DOI:** 10.1101/2024.07.09.602634

**Authors:** Tzu-Yin Chou, Yu-Zhen Ye, Chia-Jung Lien, Jui-Chun Wang, Chen-Hsuan Yang, Yen-Ting Chen, Pei-An Chao, Jian-Da Lin

## Abstract

Atherosclerosis is a growing concern in developed nations, necessitating the identification of therapeutic targets for advancing personalized medicine. Serum amyloid A3 (*Saa3*) has been linked to accelerated plaque progression by affecting cholesterol metabolism and modulation of inflammation. We hypothesize that knocking out *Saa3* (*Saa3^−/−^*) could mitigate plaque development by regulating aortic immune cell compositions during atherosclerosis progression. Using a murine model, we induced atherosclerosis via a gain-of-function mutant PCSK9-encoding adeno-associated viral vector (AAVmPCSK9) in female wild-type (WT) and *Saa3^−/−^* mice. *Saa3^−/−^* mice developed smaller plaques than WT mice, and single-cell RNA sequencing revealed significant differences in aortic immune cell populations, particularly among aortic macrophages. Aortic macrophages in atherosclerotic *Saa3^−/−^*mice represent an anti-inflammatory and tissue-repairing phenotype and the Trem2^hi^ macrophages, characterized by high *Gpnmb*, *Lpl*, and *Spp1* expressions, predominated over the typical foamy macrophages in *Saa3^−/−^* compared to WT mice. Notably, SAA3 regulates cholesterol metabolism and inflammatory responses in foamy macrophages. Our study highlights *Saa3* as a key modulator of aortic immune cells that impact atherosclerosis progression.

## Introduction

Atherosclerosis, a major cardiovascular concern in developed countries [1], is a chronic arterial disease characterized by plaque formation within arterial walls. This process involves complex interplays among various cellular components and inflammatory mediators that drive inflammation and lipid accumulation. Atherosclerosis initiation involves low-density lipoprotein (LDL) accumulation in the coronary artery intima, which upon oxidation to ox-LDL by risk factors, activates nuclear factor kappa B (NF-κB) [2, 3]. This leads to the expression of adhesion molecules like vascular cell adhesion molecule-1 (VCAM-1) and intercellular adhesion molecule-1 (ICAM-1) on the endothelium, aiding monocyte recruitment into the vascular wall [4]. These recruited monocytes then convert into macrophages under the influence of macrophage colony-stimulating factor (M-CSF), and release inflammatory cytokines such as interleukin (IL)-1β, IL-12, and tumor necrosis factor (TNF)-α, escalating the inflammatory response [5]. As the disease progresses, these macrophages become foam cells by uptaking ox-LDL, which is crucial for plaque formation and intimal thickening.

Macrophages are the predominant type of immune cell found in aortic plaques but display a wide range of phenotypes and functions. At the onset of atherosclerosis, macrophages ingest ox-LDL via receptors like CD36 and SR-A, forming cholesterol-rich foam cells. Continued lipid intake without effective cholesterol removal leads to foam cell death, creating a necrotic core in advanced plaques [6, 7]. Single-cell RNA sequencing (scRNA-seq) of monocyte-derived macrophages has revealed that macrophage heterogeneity is beyond M1/M2 definitions and associated with different states of atherosclerosis, where progression is linked to more complex states than regression [8]. This technique has revealed various macrophage subsets within atherosclerotic plaques, each characterized by unique markers and roles contributing to aortic plaque progression or regression. Notably, three main groups are identified: tissue-resident-like macrophages, marked by *Folr2* and *Lyve1*, which promote tissue stability; inflammatory macrophages, with markers such as *Tnf* and *Il1b*, which activate immune responses; and anti-inflammatory TREM2^hi^ macrophages, associated with cholesterol metabolism and identified by markers such as *Trem2* [9, 10]. This diversity in macrophage functions underscores the complexity of atherosclerosis and offers multiple targets for therapy.

The Serum Amyloid A (SAA) protein family, including SAA3, represents a group of acute-phase proteins central to immune responses and inflammatory processes. Unlike humans, who lack a functional SAA3 protein due to a premature stop codon and resulting frameshift mutation that renders the SAA3 gene a pseudogene [11], mice express a functional SAA3 and mouse SAA3 shares 70% amino acid homology with human SAA1 [12]. Its tissue distribution and expression patterns, particularly in adipose tissue, align more closely with human SAA1 than with mouse SAA1 or SAA2, suggesting a functional similarity between mouse SAA3 and human SAA1 [12]. Furthermore, sex-dimorphisms have been reported in the context of atherosclerosis, SAA3-associated inflammatory responses, and microbiota-mediated regulation of immune and metabolic processes. Smit et al. (2024) found that in aged atherosclerotic *Ldlr^−/−^* mice, plaque volume was similar between males and females, but female mice presented plaques with higher collagen and cholesterol crystal content, along with a smaller necrotic core and fewer macrophages compared to males [13]. *Saa3* has been shown to have sexually dimorphic effects on atherosclerosis, with deficiency being atheroprotective in male mice but being proatherogenic in female mice [14]. This response is suggested to be influenced by sex steroids, as 17β-estradiol promotes a pro-inflammatory profile in male *Saa3^−/−^* macrophages [14].

However, another study indicates a significant upregulation of SAA3, but not SAA1 or SAA2, in the adipose tissue of conventional mice, compared to germ-free mice [15]. The presence of intestinal microbiota significantly upregulated SAA3 expression in adipose tissue and colonic tissue, with evidence indicating that its colonic regulation is partially mediated through the Toll-like receptor (TLR)/MyD88/NF-κB signaling axis [12]. These data suggest that *Saa3*-associated sex-specific differences in atherosclerosis progression and inflammatory responses may result from its interaction with confounding factors, such as the microbiota, which modulate SAA3 levels and sex dimorphism in atherosclerosis.

Previous studies suggest that mouse *Saa3* modulates local injury and inflammatory responses, as its mRNA and protein expression in macrophages increases with the dose of lipopolysaccharide (LPS) [12]. It also indicates that increased SAA expression correlates with the aortic plaque progression in *ApoE^−/−^* mice, accompanied by elevated levels of IL-6 and TNF-α. This upregulation potentially facilitates the infiltration of M1 macrophages into atherosclerotic regions, thereby promoting atherosclerosis progression [16]. Mouse *Saa3* is actively expressed, predominantly in macrophages and adipose tissues [17], and the activation of the *Saa3* promoter is regulated through C/EBPβ signaling, which in turn activates macrophages, underscoring the complex interaction within adipose tissue [18]. Pulmonary exposure to nanoparticles induces pronounced acute phase responses, with mouse *Saa3* identified as the most differentially regulated gene, demonstrating prolonged upregulation in lung tissue and correlating with neutrophil infiltration and systemic SAA3 protein levels [19]. Notably, SAA3 has been reported to promote atherosclerosis, and overexpressed SAA3 increases atherosclerotic plaque size [20]. Overall, these studies support the role of SAA3 in regulating macrophage function during atherosclerosis, though the specific type of aortic macrophages that regulate aortic plaque progression and the underlying mechanisms remain elusive.

To assess how *Saa3* deficiency impacts aortic immune cell composition and functionalities in contributing to atherosclerosis progression, we applied scRNA-seq using the BD Rhapsody platform on immune cells from aortic arches of female *Saa3* knockout (*Saa3^−/−^*) mice and littermate wild-type (WT) control mice during atherosclerosis. Our investigation seeks to delineate the impact of *Saa3* deficiency on the phenotypic and functional landscape of aortic macrophages and other immune cell populations. By comparing the immune cell profiles between *Saa3^−/−^*and WT mice, we aim to uncover the molecular mechanisms by which SAA3 influences macrophage or other immune cell types-mediated aortic plaque progression.

## Results

### *Saa3* deletion reduced aortic plaque progression but increased peripheral lipid levels

To determine the phenotype of atherogenesis after *Saa3* deficiency, we utilized a C57BL/6 wild-type (WT) mouse-based atherosclerosis model independent of *Ldlr*^−/−^ mice breeding [8, 21, 22]. Plaque progression was initiated by injecting the mice with an adeno-associated viral vector expressing a gain-of-function mutant of protein convertase subtilisin/kexin type 9 (AAVmPCSK9), resulting in LDL receptor deficiency and hypercholesterolemia. Both female C57BL/6 WT and C57BL/6 *Saa3^−/−^*mice were treated with AAVmPCSK9 and fed the Western diet (WD) for 18 weeks to induce hypercholesterolemia and aortic plaque formation. We longitudinally collected plasma from mice at 0, 2, 4, 8, 12, and 16 weeks on WD, and the immune profiling was performed on sorted CD45+ cells from aortic arches by single-cell RNA-seq at 18 weeks on WD (Fig. 1a). We first determined total cholesterol levels in the plasma of WT and *Saa3^−/−^*mice and a baseline comparison at weeks 0 revealed no significant differences, with levels below 100 mg/dL (Fig. 1b). Over time, *Saa3^−/−^*mice developed higher cholesterol levels than WT, with a slight increase at weeks 2 through 12, and a significant rise by weeks 16 (Fig. 1b).

**Fig. 1.**
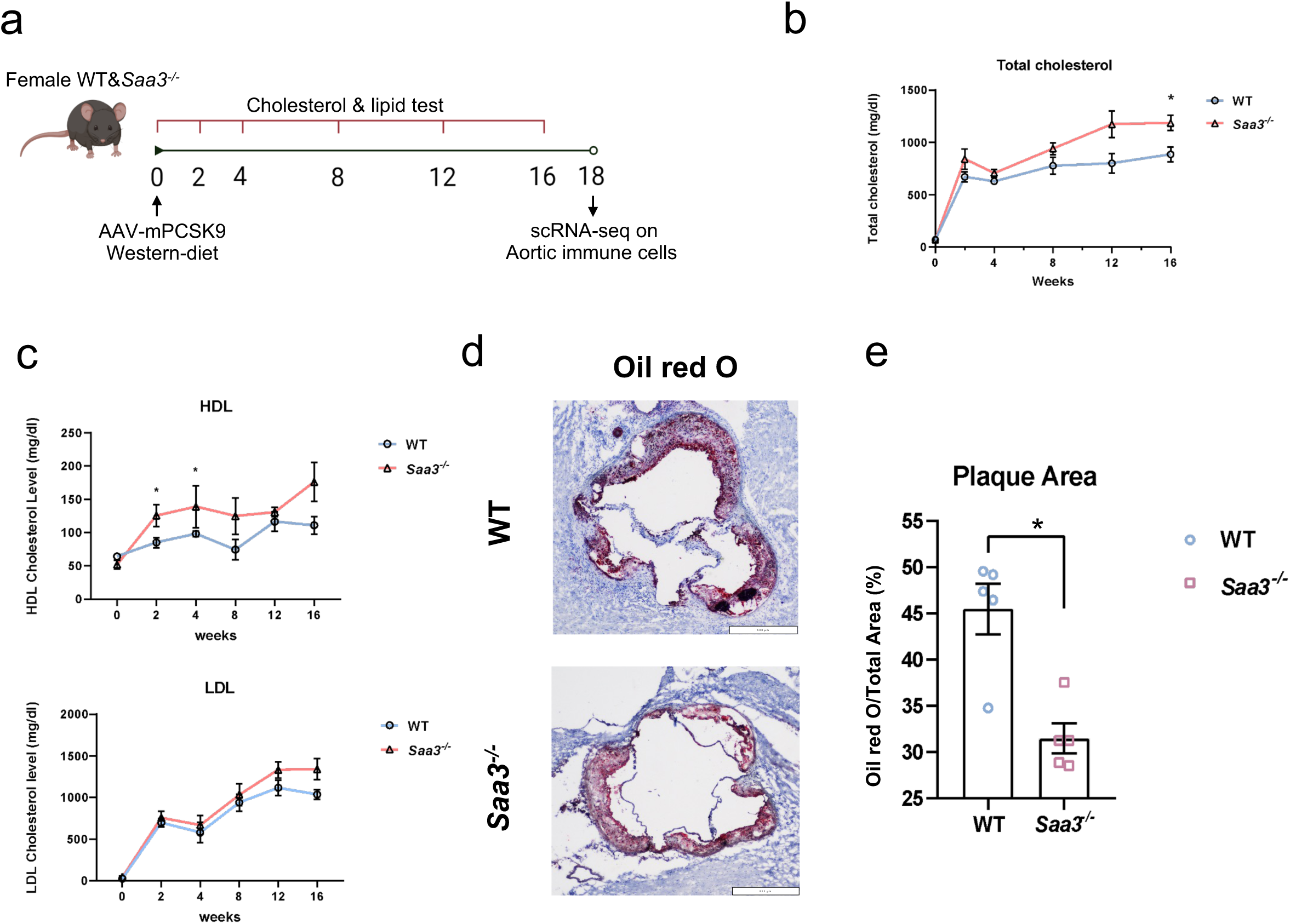
Effects of *Saa3* deletion on peripheral lipid profiles and aortic plaque progression during atherosclerosis. (**a**) Schematic of an experimental model for the time points of plasma collections and single-cell RNA sequencing (scRNA-seq) on aortic immune cells. C57BL/6 wild-type (WT) and *Saa3^−/−^* female mice were injected with AAVmPCSK9 and fed with the Western diet for 18 weeks. Then aortic arches were digested to single-cell suspensions and FACS-sorted aortic CD45+ cells were subjected to scRNA-seq on the BD Rhapsody platform from WT and *Saa3^−/−^* mice. (**b**) Plasma total cholesterol levels in WT and *Saa3^−/−^*mice subjected with ssAAV8/hAAT-mPCSK9 and fed with the Western diet for 0, 4, 8, 12, and 16 weeks. (**c**) Measurement of high-density lipoprotein (HDL) cholesterol levels and low-density lipoprotein (LDL) cholesterol levels in plasma from WT and *Saa3^−/−^* mice over a period of 0, 2, 4, 8, 12, and 16 weeks. (**d**) Representative Oil Red O staining of aortic root sections in WT and *Saa3^−/−^* mice at 18 weeks post Western diet. Scale bar: 100 μm. (**e**) As in (d), quantification of the lesion area of aortic root sections stained with Oil Red O. All data are presented as mean ± SEM; Statistically significant differences between the two groups at respective time points were denoted by asterisks (\**p* < 0.05), determined using an unpaired Mann-Whitney U-test. Each symbol represents an individual mouse; WT mice (n=5), *Saa3^−/−^* mice (n=5).

Research demonstrates that serum amyloid A (SAA) attenuates the anti-inflammatory and antioxidative functions of high-density lipoprotein (HDL) by facilitating its binding to adipocyte and macrophage-derived glycoproteins [23]. We conducted high-density lipoprotein (HDL)/ low-density lipoprotein LDL analyses that revealed an increase in peripheral HDL levels during the initial phase of atherosclerosis progression (Fig. 1c). Not only did HDL levels demonstrate a rising trend with statistically significant increases in the *Saa3^−/−^*group compared to WT at weeks 2 and 4, but the absence of *Saa3* also correlated with a potential increase in LDL levels in the bloodstream (Fig. 1c).

Overexpression or suppression of SAA3 can increase or decrease aortic lesion area in *ApoE^−/−^* mice [20]. We also determined aortic plaque sizes between WT and *Saa3^−/−^* mice by histologically evaluating aortic root sections stained with Oil Red O (Fig. 1d). It demonstrated a significant decrease in plaque area in *Saa3^−/−^* compared to WT mice, with mean values of 31.50% and 45.48%, respectively (Fig. 1e). These observations demonstrate that *Saa3* deficiency increases peripheral lipid levels but attenuates aortic plaque formation, suggesting the complex roles of *Saa3* in modulating lipid metabolism and atherogenesis.

### *Saa3* deficiency alters intercellular immune communications in aortic arches during atherosclerosis progression

All immune cells express CD45, a major transmembrane glycoprotein expressed on all hematopoietic cells [24] and some cardiac fibroblasts [25]. To elucidate the immunological impact of *Saa3* deficiency within the aortic arch and its putative role in the pathogenesis of atherosclerosis, we performed a single-cell RNA sequencing on FACS-purified CD45+ cells (Supplementary Fig. 1). After data processing, possible contaminated cell removement, anchor-based dataset integration [26], and cell clusters visualized by Uniform Manifold Approximation and Projection (UMAP), it revealed the immune cell compositional variances in aortic arches between female WT and *Saa3^−/−^*mice during atherosclerosis progression (Fig. 2). Utilizing the cell annotation tool by *SingleR* [27], we classified the cell phenotypes, enabling initial taxonomical segregation within the distinct clusters (Fig. 2a). Our analysis identified 9 cell subpopulations, encompassing B cells, dendritic cells (DCs), innate lymphoid cells (ILCs), macrophages, monocytes, neutrophils, natural killer T (NKT) cells, T cells, and ψ8 T cells (Tgd) (Fig. 2a). Despite similar UMAP topologies between WT and *Saa3^−/−^* aortic cell populations (Fig. 2a), *Saa3^−/−^* mice showed increased T cells, B cells, monocytes, ψ8 T cell, and neutrophils proportions but decreased macrophages, innate lymphoid cells (ILC), and NKT cells compared to WT mice (Fig. 2b).

**Fig. 2.**
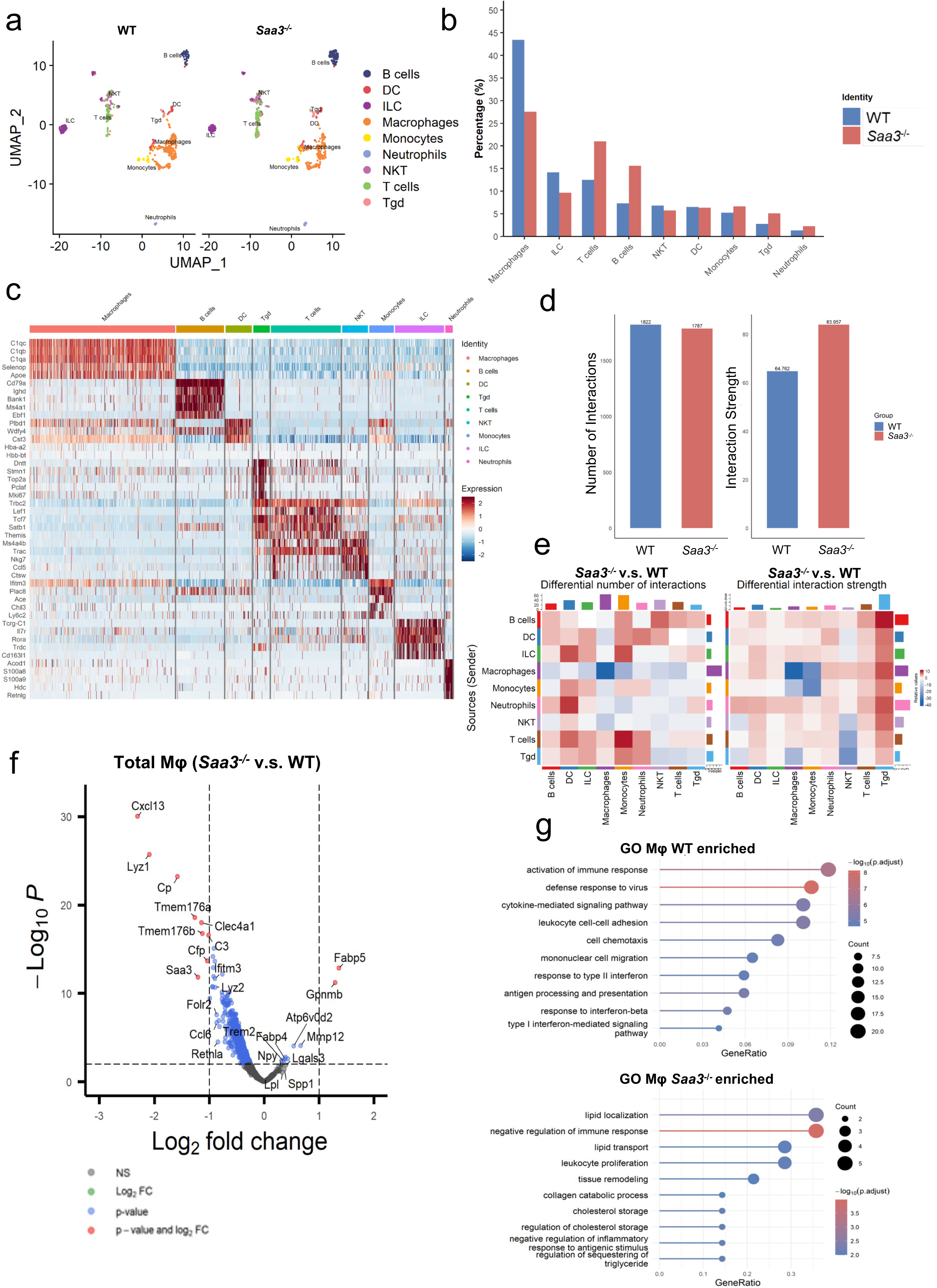
S*a*a3 deficiency alters immune intercellular communication in aortic arches during atherosclerosis progression. **(a)** UMAP visualization compares the distribution of cell types in aortic arches between WT and *Saa3^−/−^* mice. Unbiased cell type annotations against the IMMGEN database are colored and labeled accordingly on the plot. **(b)** Cell type composition by percentages of WT and *Saa3^−/−^* mice (blue, WT; red, *Saa3^−/−^*) in total sorted CD45+ cells. **(c)** Heatmap of top 5 marker genes of each cell type in aortic arches from WT and *Saa3^−/−^* mice. The selection of these top marker genes was based on the results of a Wilcoxon test for differential expression. **(d)** Bar plots show the number (left) and strength (right) of inferred interactions of intercellular communication networks in aortic arches from WT and *Saa3^−/−^* mice. **(e)** Heatmap visualization of intercellular communication networks in aortic arches from WT and *Saa3^−/−^* mice. The red and blue colors represent increased or decreased signaling in the aortic arches of *Saa3^−/−^* mice compared to the WT mice. The top-colored bar plot represents the sum of the columns of values (incoming signaling), and the right-colored bar plot represents the sum of a row of values (outgoing signaling). **(f)** Volcano plot illustrating differential gene expression in total macrophages within aortic arches of *Saa3^−/−^*versus WT mice. The x-axis displays the log_2_ fold change, while the y-axis shows the –Log_10_ *p*-value. Grey dots denote genes without significant differences, green dots indicate genes with significant fold changes, and red dots mark genes significant in both fold change and *p*-value. The dashed vertical lines represent the fold change boundaries, and the dashed horizontal line demonstrates the p-value significance threshold. **(g)** Top 10 biological processes enriched in total macrophages within the aortic arches of WT mice compared to *Saa3^−/−^* mice (upper panel) or *Saa3^−/−^* mice compared to wild-type mice (lower panel). Enrichment analysis utilized GO pathways to identify significant biological processes. The gene ratio is plotted along the x-axis, and the enriched biological processes are listed on the y-axis. A color gradient has been applied to indicate the –log_10_ adjusted *p*-value (p.adjust) of each term, with the size of each point reflecting the number of genes associated with the respective pathway.

Heatmaps of differential gene expression profiles supported these *SingleR*-annotated cell type differences, highlighting the top 5 marker genes per subset (Fig. 2c). *Cellchat* has been used to infer, analyze, and visualize cell-cell communication networks in scRNA-seq datasets [27]. To dissect the variances in signaling dynamics in aortic cell types between WT and *Saa3^−/−^*mice, we employed *CellChat* to evaluate the intercellular communication networks and assess the number and intensity of cellular interactions within the respective cohorts (Figs. 2d-2e). The resultant metrics demonstrated 1,822 and 1,787 interactions across all cell types in aortic arches between WT and *Saa3^−/−^* mice, with interaction strength for *Saa3^−/−^* cell types quantified at 83.957 compared to 64.762 for WT (Fig. 2d). To rigorously dissect the discrete intercellular communication patterns responsible for the observed differential signaling, we interrogate the heatmaps that delineate both the frequency and magnitude of interaction strengths across aortic cell types (Fig. 2e). Increased magnitude of intercellular interaction strengths was observed in the aortic arches of *Saa3^−/−^* mice compared to their WT mice (Fig. 2d). The increased interaction magnitudes were primarily directed from most immune cells toward γδ T cells (Tgd), while reduced interactions were directed from macrophages toward other immune cells in *Saa3^−/−^* mice compared to WT mice (Fig. 2e). These patterns suggest that the absence of the *Saa3* gene exerts a profound regulatory influence on the intercellular communication network among immune cells, potentially revealing critical insights into the regulatory mechanisms of immune function within the aortic niche.

### *Saa3* depletion increases biological processes in aortic macrophages that are highly associated with lipid metabolism and inflammatory recruitment during atherosclerosis

*CellChat* analysis revealed significant differences in cell-cell interaction signals between aortic macrophages and other aortic immune cell populations within the *Saa3^−/−^* and WT group (Figs. 2d-2e). To delve deeper, we assessed the transcriptomic variances in macrophages from *Saa3^−/−^* versus WT mice. Our volcano plot delineated distinct gene expression profiles, with genes such as *Fabp5* and *Gpnmb* are upregulated in *Saa3^−/−^*mice, exceeding the established log_2_ fold change threshold of >1 and p-values < 0.01 (Fig. 2f). *Fabp5*, which plays a central role in the intracellular transport of lipids, is integral to the uptake, transport, and metabolism of fatty acids [28]. *Gpnmb* has been shown to protect against obesity-related metabolic disorders by mitigating macrophage inflammatory capacity [29]. Conversely, genes such as *Cxcl13*, *Lyz1*, *Tmem176a*, *Clec4a1*, *C3*, and *Cfp* are downregulated in *Saa3^−/−^* macrophages (Fig. 2f). Studies have demonstrated that a lack of *Cxcl13* correlates with a diminished presence of atherosclerotic conditions, likely due to lower recruiting inflammatory granulocytes within the lesions [30] [31]. *C3* expression in macrophages has been linked to exacerbating atherosclerosis by limiting macrophage efferocytosis, contributing to plaque development and inflammation [32]. *Cfp* has been linked to regulating complement activation and orchestrating iron homeostasis [33]. Gene set enrichment analysis, conducted using the *ClusterProfiler* package [34], revealed distinct gene ontology (GO) pathways in aortic macrophages (Fig. 2g). WT aortic macrophages were enriched for pathways associated with enhanced immune cell recruitment and chemotaxis, whereas *Saa3^−/−^* aortic macrophages exhibited enrichment for pathways related to lipid localization and transport (Fig. 2g, Supplementary Table 1). Overall, these data suggest the phenotypes of aortic macrophages were impacted by reduced recruitment of inflammatory macrophages, which helped mitigate inflammation potentially influencing lesion formation and stability.

### Altered macrophage compositions and enhanced Trem2^hi^ macrophage presence linked to diminished aortic plaque progression in *Saa3^−/−^* mice

Macrophage (Mφ) polarization, along with several distinct clusters, has been identified as increasingly prevalent during the atherosclerosis progression than regression [8, 35]. To dissect the functional heterogeneity of aortic macrophage populations in WT and *Saa3^−/−^* mice during atherosclerosis, we classified macrophages into subpopulations and compared the top differentially expressed genes from each cluster against our previously published dataset, where we profiled single-cell sorted *Cx3cr1*-derived aortic macrophages from lineage-tracing mice during atherosclerosis progression and regression (hereinafter referred to as “*Lin regression study*”)[8]. There are five distinct macrophage clusters in the aortic arches of WT and *Saa3^−/−^* mice during atherosclerosis (Fig. 3a). CD74^hi^MHCII^hi^ Mφ is the most abundant macrophage subset in both genotypes, with CD74^hi^MHCII^hi^, Folr2^hi^, and Retnla^hi^Ear2^hi^ Mφ being more prevalent in WT mice compared to *Saa3^−/−^* mice (Fig. 3b). Conversely, Trem2^hi^ and Apbb2^hi^ Mφ are found in higher percentages within the aortic arches of *Saa3^−/−^*mice than WT mice (Fig. 3b).

**Fig. 3.**
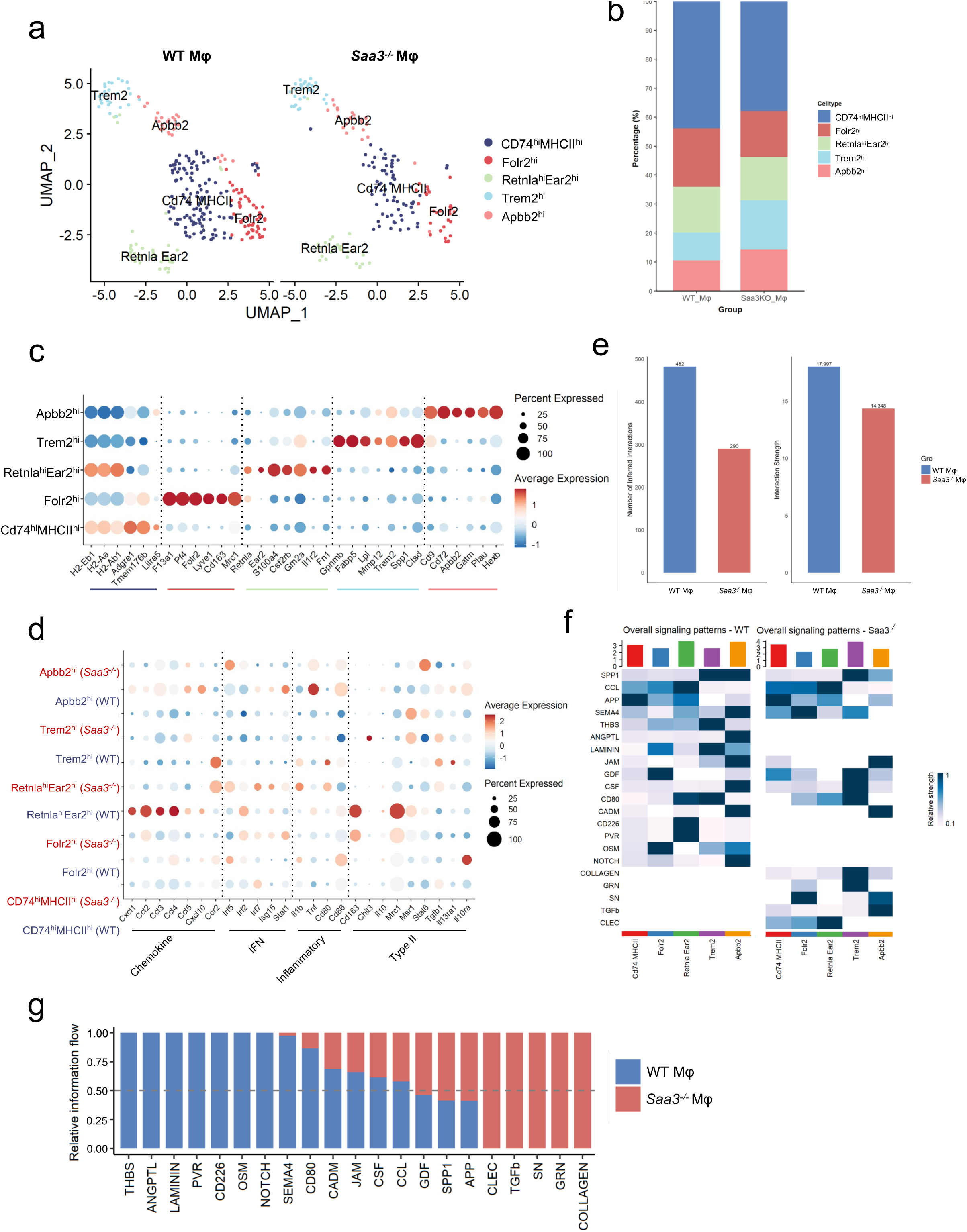
Delineation of macrophage sub-cluster in aortic arches between WT and *Saa3^−/−^* mice during atherosclerosis progression. (**a**) UMAP plots show the sub-clusters of macrophages (Mφ) in aortic arches between WT (left) and *Saa3^−/−^*(right) mice. Macrophages are delineated based on their gene expression profiles: CD74^hi^MHCII^hi^ Mφ in navy blue, Folr2^hi^ Mφ in red, Retnla^hi^Ear2^hi^ Mφ in green, Trem2^hi^ Mφ in sky blue, and Apbb2^hi^ Mφ in pink. (**b**) The bar graph displays the percentage composition of various macrophage subpopulations within total macrophages from WT and *Saa3^−/−^* mice. Different colors represent distinct macrophage subpopulations as indicated in **(a)**. (**c**) Dot plot representing the expression profiles of five sub-macrophage populations. The x-axis lists genes with differential expression, while the y-axis indicates the sub-macrophage clusters. (**d**) Dot plots show the expression profiles of macrophage markers involved in pathways of chemokine, IFN (interferon), inflammatory, and type II responses across sub-macrophage clusters. For **(c)** and **(d)**, the size of each dot corresponds to the percentage of the indicated cell cluster expressing a particular gene marker, and the color intensity signifies the mean expression level within that cell cluster. (**e**) Bar plots show the number (left) and strength (right) of inferred intercellular communication networks in total aortic macrophages from WT and *Saa3^−/−^* mice. (**f**) Heatmap visualization of overall signaling patterns in aortic macrophages of WT and *Saa3^−/−^* mice. The top-colored bar plot represents the sum of each column of values (signaling the strength of overall pathways) across sub-clusters. The gradient blue colored bar represents the relative strength of the indicated pathway. (**g**) The bar graph displays the relative information flow of various signaling molecules in aortic macrophages of WT and *Saa3^−/−^* mice. Blue bars indicate higher relative information flow in WT macrophages, while red bars indicate higher information flow in *Saa3^−/−^*macrophages.

The Cd74^hi^MHCII^hi^ Mφ, a subset identified through the *Lin regression study* [8], is characterized by high expression of genes such as *Tmem176b*, *Lilra5*, *H2-Eb1*, and *H2-Aa* (Supplementary Fig. 2a and Table 2). Folr2^hi^ Mφ exhibit unique transcriptional profiles with *F13a1, Pf4, Folr2*, and *Lyve1* (Supplementary Fig. 2a and Table 2). These markers are indicative of a tissue-resident-like macrophage phenotype [36–39] and align with the molecular signature of the Folr2^hi^ macrophage cluster previously delineated in the *Lin regression study* [8]. Retnla^hi^Ear2^hi^ Mφ is distinguished by their elevated expression of genes such as *Retnla* and *Ear2*, which suggest an early-stage transition of monocytes to macrophages within tissues [40, 41]. Additionally, these macrophages show enrichment in genes like *Gm2a* and *Fn1*, confirming their specialized state as identified in the *Lin regression study* [8]. Trem2^hi^ Mφ is defined by a gene expression signature that mirrors the population described by *Cochain et al.* [42] and *Lin regression* [8] studies and is exemplified by heightened expression of *Ctsd, Lgals3, Spp1*, and *Lpl* (Supplementary Fig. 2a and Table 2). Apbb2^hi^ Mφ is characterized by the predominant expression of *Cd9, Cd72*, and *Hexb* genes. While sharing traits with the “Trem2^hi^ Mφ,” they are uniquely distinguished by their expression of *Apbb2* (Supplementary Fig. 2a and Table 2).

On the other hand, we identified differentially expressed genes (DEGs) across five macrophage populations and conducted gene ontology (GO) enrichment analysis on biological pathways to explore potential functions (Supplementary Fig. 2b and Table 3). Compared to other macrophage clusters, Cd74^hi^MHCII^hi^ Mφ in the aortic arches showed significant enrichment in pathways related to leukocyte-mediated immunity, positive regulation of cell activation, activation of the immune response, and immunoglobulin-mediated immune responses (Supplementary Fig. 2b and Table 3). Folr2^hi^ Mφ is distinguished by activation of immune response, ameboidal-type cell migration, cell chemotaxis, regulation of angiogenesis, and regulation of viral processes (Supplementary Fig. 2b and Table 3). Retnla^hi^Ear2^hi^ Mφ were noted for their leukocyte proliferation, mononuclear cell proliferation, positive regulation of leukocyte cell-cell adhesion, positive regulation of T cell activation, and response to type II interferon (Supplementary Fig. 2b and Table 3). Trem2^hi^ Mφ is characterized by the generation of precursor metabolites and energy, aerobic respiration, oxidative phosphorylation, purine ribonucleoside triphosphate biosynthetic process, and ATP biosynthetic process (Supplementary Fig. 2b and Table 3). These functions highlight the metabolic specialization of Trem2^hi^ Mφ in energy production and biosynthesis. Apbb2^hi^ Mφ demonstrated upregulation in myeloid cell differentiation, leukocyte migration, actin filament organization, regulation of protein-containing complex assembly, and mononuclear cell migration (Supplementary Fig. 2b and Table 3).

In *Lin regression study* [8], Cd74^hi^MHCII^hi^ Mφ and Retnla^hi^Ear2^hi^ Mφ are abundant in aortic arches of progression group and we also observed these two subsets are higher in aortic arches of WT mice compared to *Saa3^−/−^* mice (Fig. 3b). *Folr2* expression is increased in human atherosclerotic plaques compared to normal arterial tissue[43] and we also observed aortic Folr2^hi^ Mφ are higher in WT mice than *Saa3^−/−^*mice (Fig. 3b). Trem2^hi^ Mφ exhibits comparable prevalence between the progression and regression cohorts in the *Lin regression study* [8], whereas both Trem2^hi^ and Apbb2^hi^ Mφ are more abundant in the aortic arches of *Saa3^−/−^* mice compared to WT mice (Fig. 3b). Trem2^hi^ Mφ is previously identified as lipid-laden foam cells within atherosclerotic lesions, underscoring their capacity for lipid processing and phagocytosis [9, 10, 35]. They are associated with anti-inflammatory responses, which may help prevent the establishment of a proinflammatory environment within the plaque [9, 35, 44]. Similar to previous findings, Trem2^hi^ Mφ expresses high *Spp1,* a gene associated with fibrotic responses indicative of tissue repair mechanisms [45, 46] (Fig. 3c, Supplementary Fig. 2a). Additionally, this subset expresses *Gpnmb*, a gene implicated in anti-inflammatory processes [29, 47], along with *Fabp5*, which is involved in the uptake and transport of fatty acids [28] (Fig. 3c). Notably, enzymes such as *Lpl*, central to lipid metabolism[48], and *Mmp12*, a contributor to extracellular matrix remodeling [49], further characterize this population (Fig. 3c). Hence, these data indicate that *Saa3^−/−^* mice exhibit higher compositions of aortic macrophages with tissue-repairing and anti-inflammatory properties, leading to reduced aortic plaque progression.

### *Saa3* modulates aortic macrophage functionalities during atherosclerosis

Investigating the transition of macrophage phenotypes from M1-like to M2-like as a therapeutic strategy recognizes the complexity of aortic macrophage phenotypes in atherosclerosis, which extend beyond conventional M1 and M2 polarization states [8, 35]. We use typical markers for Chemokine, Interferon (IFN), Inflammatory, and Type II responses to visualize the aortic macrophage subsets from WT and *Saa3^−/−^* mice (Fig. 3d). Among the comparisons between WT and *Saa3^−/−^* aortic Mφ, pronounced differences were noted in the increased expression levels of chemokines including *Cxcl1*, *Ccl2*, *Ccl3*, and *Ccl4* in *Saa3^−/−^* than WT Folr2^hi^ Mφ (Fig. 3d).

Chemokines are signaling molecules that attract various types of immune cells to sites of infection or injury. Increased expression of chemokines by tissue-resident macrophages enhances the recruitment of neutrophils and lymphocytes, which is crucial for initiating and sustaining an effective immune response [50]. Additionally, there was a higher expression of type II response markers including *Cd163, Mrc1*, and *Msr1* [51–54] in *Saa3^−/−^*than WT Folr2^hi^ Mφ (Fig. 3d). Moreover, Il10ra showed higher expression in *Saa3^−/−^* than WT Cd74^hi^MHCII^hi^ Mφ, and Il10ra is required for protection against atherosclerosis in low-density lipoprotein receptor knockout mice [55]. Furthermore, we observed higher expression levels of interferon regulatory and downstream markers such as *Irf5*, *Irf7*, *Isg15*, and *Stat1* in WT than *Saa3^−/−^*Retnla^hi^Ear2^hi^ Mφ (Figure 3d), which may indicate an enhanced interferon response in these cells [56, 57]. These expression patterns indicate a significant alteration in gene expression profiles and macrophage composition between the two genotypes, suggesting that deletion of the *Saa3* may modulate macrophage recruitment, type 2 immune responses, and inflammatory/anti-inflammatory phenotypes.

### *Saa3* modulates aortic macrophage communications that impact inflammatory processes

To determine the role of *Saa3* in aortic macrophage communication during atherosclerosis, we also performed a comparative analysis of cell interactions using *CellChat* [27] in both WT and *Saa3^−/−^* macrophages. Our findings indicate a substantial decline in intercellular communication in *Saa3^−/−^*macrophages, with interaction events decreasing from 482 in WT to 290 in *Saa3^−/−^*cells (Fig. 3e).

Correspondingly, we observed a significant drop in the strength of these interactions, with the average interaction strength in WT macrophages being 17.997 compared to a lower average of 14.348 in *Saa3^−/−^* macrophages, mirroring the decrease in the number of interaction events (Fig. 3e).

The heatmap depicts the overall signaling patterns observed in various macrophage subpopulations from WT and *Saa3^−/−^* mice (Fig. 3f), and the bar graph shows the relative information flow of these signaling pathways between WT and *Saa3^−/−^* macrophages (Fig. 3g). In the context of inflammatory-mediated atherosclerosis progression, WT macrophages present higher signaling activity in THBS, ANGPTL, LAMININ, PVR, CD226, OSM, NOTCH, CADAM, and JAM pathways compared to *Saa3^−/−^*macrophages (Figs. 3f-3g). In THBS signaling, *Thbs1* is a critical regulator of macrophage interleukin-1β (IL-1β) production and activation via CD47, highlighting the complex interplay between thrombospondins and immune modulation [58]. In ANGPTL signaling, ANGPTL2 promotes vascular inflammation and accelerates atherosclerosis progression, with increased cardiac ANGPTL2 production linked to heart failure and pathological cardiac remodeling by influencing inflammatory responses and affecting both cardiomyocytes and macrophages[59, 60]. In LAMININ and JAM signaling, LAMININ and junctional adhesion molecules (JAM) are involved in monocyte-to-macrophage differentiation and macrophage migration, playing roles in immune cell dynamics and atherosclerotic disease modulation [61, 62]. PVR significantly contributes to atherosclerosis by facilitating leukocyte migration across the endothelium, thereby promoting inflammation [63]. When loss of CD226, a costimulatory molecule, has been observed to protect against atherosclerosis by preventing foam cell formation and reducing plaque buildup [64].

Oncostatin M (OSM) signaling promotes atherogenesis and plaque vulnerability by promoting the differentiation of vascular smooth muscle cells into osteoblastic phenotypes via M1 macrophages, potentially leading to plaque instability [35].

NOTCH signaling plays a significant role in atherosclerosis development, with *Notch-1* deletion inducing anti-inflammatory M2 effects through the PI3K-Akt pathway, contrasting with its increased presence in vulnerable human plaques [65].

In contrast to their WT counterparts, *Saa3^−/−^* macrophages exhibit elevated expression of signaling molecules such as COLLAGEN, GRN, SN, TGFb, and CLEC. Collagen plays a crucial role in modulating macrophage functions and improving plaque stability, with an increased presence in M2-polarizing macrophages within the plaque [66, 67]. GRN exerts anti-atherogenic effects by enhancing nitric oxide production in endothelial cells through the Akt/eNOS signaling pathway, forming a complex with apolipoprotein A-I to stabilize atherosclerotic plaques. In addition, in atherosclerotic mice, its deficiency leads to increased expression of adhesion molecules and inflammatory cytokines [68–70]. TGF-β signaling contributes to the reduction and stabilization of atherosclerotic plaques by altering macrophage functions, and its overexpression is linked to smaller lesions, increased collagen deposition, smooth muscle cell infiltration, and reduced macrophage presence, ultimately fostering a more stable lesion phenotype [71]. CLEC signaling, specifically through Clec4a2, a C-type lectin receptor, protects against atherosclerosis by promoting homeostatic functions within the vessel wall, enhancing cholesterol metabolism, and limiting immune responses to toll-like receptors [72]. In summary, this analysis confirms that aortic macrophages in WT mice are more associated with inflammatory and migration processes, whereas aortic macrophages in *Saa3^−/−^* mice are more related to vascular protection and aortic plaque stabilization.

### Differential effects of *Saa3* deficiency on macrophage subpopulations and ligand-receptor interactions

We further investigated the differential effects of *Saa3* deficiency on five macrophage subpopulations. Our observations revealed that, aside from Trem2^hi^ Mφ, the gene expression profiles of CD74^hi^MHCII^hi^ Mφ, Folr2^hi^ Mφ, Retnla^hi^Ear2^hi^ Mφ, and Apbb2^hi^ Mφ in *Saa3^−/−^* mice exhibited substantial downregulation of numerous genes when compared to their WT counterparts (Figs. 4a-4d). We also performed GO term enrichment pathway analysis on the downregulated DEGs in these four *Saa3^−/−^* macrophage subpopulations compared to WT (Figs. 4e-4h). The analysis revealed common downregulated genes in *Saa3^−/−^* mice, including *Lyz1*, *Lyz2*, other lysozyme-related genes, *Cxcl13*, and *Ccl8*, which are associated with pathways involved in cell killing and the destruction of other organisms, suggesting that the absence of *Saa3* impairs macrophage functions in combating pathogens as previous studies [73, 74] (Figs. 4e-4h). Specifically, Folr2^hi^ Mφ in *Saa3^−/−^* mice showed significant downregulation of *Cxcl13* (Fig. 4b). Previous study indicates patients with carotid atherosclerosis exhibit elevated plasma levels of CXCL13, especially in symptomatic cases, and higher CXCL13 expression within atherosclerotic carotid plaques compared to non-atherosclerotic vessels [75]. CCL8 is a potent chemoattractant for monocytes and macrophages, directing them to areas of inflammation and infection [76, 77]. Retnla^hi^Ear2^hi^ Mφ in WT mice show higher enrichment in ERK1 and ERK2 cascade, positive regulation of angiogenesis, and lipid storage (Figs. 4c, 4g). ERK1 and ERK2 signaling pathways regulate macrophage function in atherosclerosis.

**Fig. 4.**
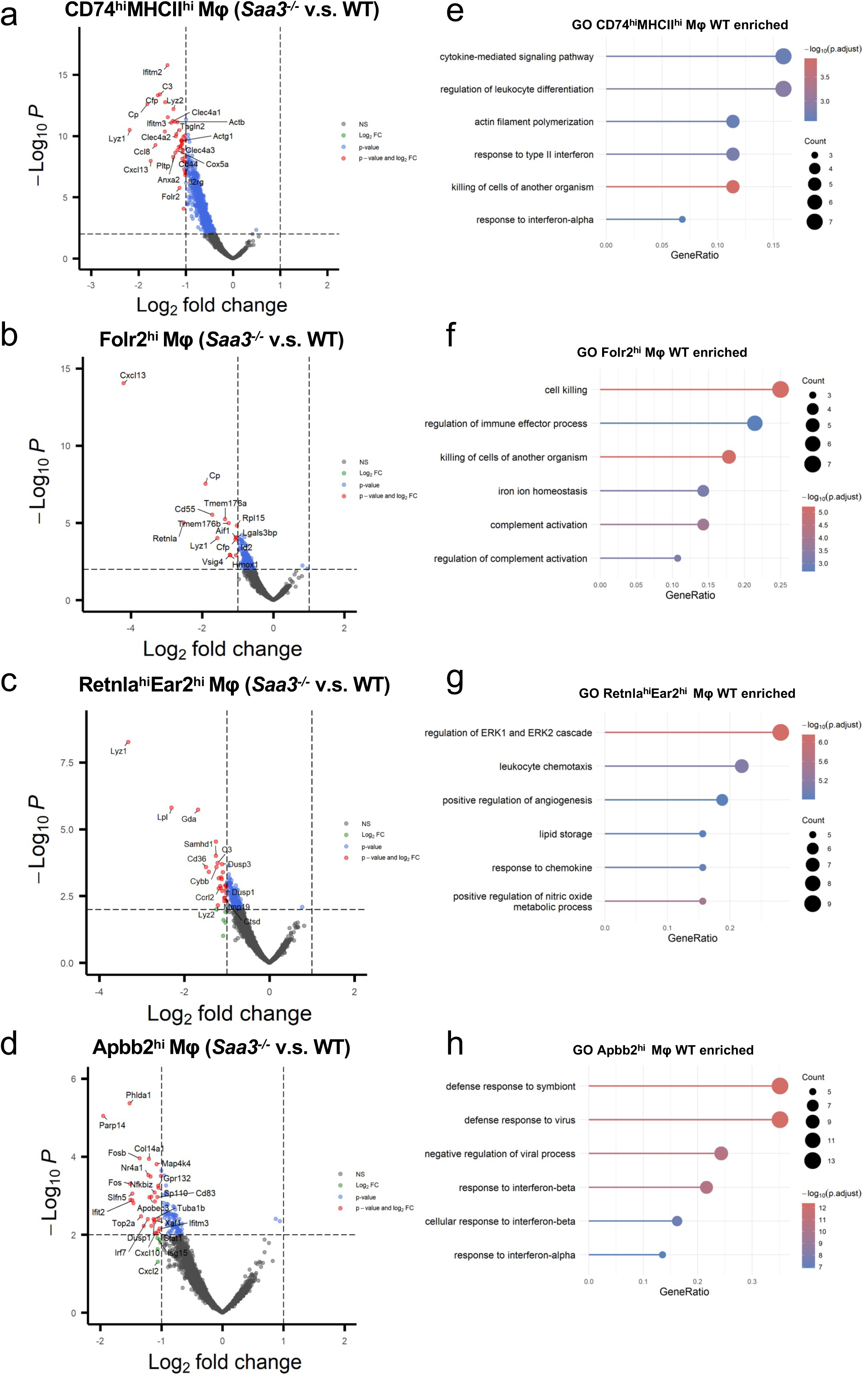
*Sa*a3 deficient impacts gene expression profiles of aortic macrophages. **(a)-(d)** Volcano plot illustrating differential gene expression in four sub-macrophage populations (CD74^hi^ MHCII^hi^, Folr2^hi^, Retnla^hi^ Ear2^hi^, and Apbb2^hi^ macrophage(Mφ)) within aortic arches of WT versus *Saa3^−/−^* mice. The x-axis displays the log_2_ fold change, while the y-axis shows the –Log_10_ p-value. Grey dots denote genes without significant differences, green dots indicate genes with significant fold changes, and red dots mark genes significant in both fold change and *p*-value. The dashed vertical lines represent the fold change boundaries, and the dashed horizontal line demonstrates the *p*-value significance threshold. **(e)-(h)** The bubble plot presents the GO enrichment analysis for CD74^hi^ MHCII^hi^, Folr2^hi^, Retnla^hi^ Ear2^hi^, and Apbb2^hi^ aortic macrophage (Mφ) from WT and *Saa3^−/−^* mice. The x-axis represents the GeneRatio. The y-axis lists the enriched GO terms. Bubble size corresponds to the count of DEGs associated with each GO term. The color gradient from blue to red represents the –log_10_ (adjusted *p*-value) of the enrichment.

Deletion of *Erk1* reduces macrophage migration and plaque lipid content in LDL receptor-deficient mice on a high-fat diet [78, 79]. Apbb2^hi^ Mφ in WT mice enrich in defense response to virus and interferon (Figs. 4d, 4h). This may indicate that Apbb2^hi^ Mφ in WT mice has more inflammatory response features than in *Saa3^−/−^* mice, contributing to chronic inflammation, plaque instability, and progression [80]. To elucidate the specific ligand-receptor interactions contributing to the observed differences in pathway patterns between *Saa3^−/−^* and WT groups, we conducted a comparative analysis of ligand-receptor signaling within five subpopulations of macrophages (Supplementary Figs 3a-3e). The analysis encompassed an assessment of the differential expression of ligands and receptors and the computation of interaction probabilities, thereby enabling the dissection of signaling discrepancies between the two genotypic conditions. Since aortic Trem2^hi^ Mφs are more abundant in *Saa3^−/−^* mice, we focus on ligand-receptor interactions within Trem2^hi^ Mφ under both WT and *Saa3^−/−^* conditions (Supplementary Fig. 3d). Prominently, pairs such as Spp1-Cd44 and Spp1-integrins (Itgav+Itgb1, Itga5+Itgb1, Itga4+Itgb1), exhibit a heightened probability of interaction in the signal transduction pathways of Trem2^hi^ Mφ within the *Saa3^−/−^* mice (Supplementary Fig. 3d). Spp1 is established to bind integrins Itgav, Itgb1, Itgb5, and Itga4 (Col4a2-Sdc44, Col4a2-Cd44, Col4a2-(Itga9+Itgb1), Col4a1-Sdc44, Col4a1-Cd44, Col4a1-(Itga9+Itgb1)), influencing cellular adhesion and migration crucially [81, 82] (Supplementary Fig. 3d). Furthermore, interactions involving Col4a2 and Col4a1 with their partners are exclusively observed in the *Saa3^−/−^* condition (Supplementary Fig. 3d). As major constituents of the basement membrane, type IV collagen may have implications for the stability of atherosclerotic plaques [83]. In contrast, the CCL-CCR signaling axis, encompassing Ccl9-Ccr1, Ccl8-Ccr1, Ccl7-Ccr1, Ccl6-Ccr2, Ccl6-Ccr1, Ccl3-Ccr5, and Ccl3-Ccr1, is exclusively active in the WT condition within Trem2^hi^ Mφ transitioning to Retnla^hi^Ear2^hi^ Mφ (Supplementary Fig. 3d). These chemokine-mediated communications facilitate the recruitment of macrophages or monocytes to sites of inflammation, contributing to the formation and progression of atherosclerotic lesions and plaque vulnerability [84, 85]. Parallel observations were noted for the ligand-receptor pairs Cd74^hi^MHCII^hi^ Mφ, Folr2^hi^ Mφ, Retnla^hi^Ear2^hi^ Mφ, and Apbb2^hi^ Mφ to Retnla^hi^Ear2^hi^ Mφ (Supplementary Figs. 3a-3e). It is posited that these altered signaling patterns may underlie the observed reductions in plaque burden and macrophage recruitments in *Saa3^−/−^* mice.

### Trem2^hi^ Mφ potentially mitigates aortic plaque development by enhancing lipid uptake, promoting an anti-inflammatory environment, and increasing tissue remodeling

In atherosclerosis, *Saa3* deletion precipitates substantial changes in macrophage gene expression profiles, within the Trem2^hi^ Mφ, as evidenced by differential expression between WT and *Saa3^−/−^* groups (Fig. 5a). The upregulation of genes in *Saa3^−/−^* than WT Trem2^hi^ Mφ such as *Fabp4, Fabp5,* and *Lpl*, which are central to lipid metabolism [86, 87], as well as *Ctsk*, has been implicated in promoting disturbed flow-induced endothelial inflammation [88]. Other genes including *Gpnmb, Atp6v0d2, Mrc1, Mmp12*, and *Spp1* also exhibit altered expression (Fig. 5a). Studies suggest that *Gpnmb* may be anti-inflammatory by promoting inflammation resolution, as it can attenuate pro-inflammatory responses and promote M2 polarization in macrophages [47, 89, 90]. The suppression of *Gpnmb* has been associated with enhanced atherosclerotic plaque formation, whereas its overexpression correlates with decreased plaque development [91]. Augmented *Atp6v0d2* expression activates autophagy as a compensatory response to hemodynamic pressure, which indicates its role in maintaining plaque stability and potentially curtailing the progression of atherosclerosis [92, 93]. *Mrc1* appears to modulate macrophage polarization toward an anti-inflammatory phenotype [94], while *Mmp12* plays a pivotal role in attenuating inflammation and fostering resolution via targeted proteolysis of chemokines [95, 96]. Spp1+ foamy macrophages have been linked to signaling pathways that might abate plaque progression and stabilize susceptible lesions [97]. These genes correspond to GO enrichment pathways such as lipid localization, fatty acid transport, and the positive regulation of macrophage-derived foam cell differentiation (Fig. 5b). These results suggest that *Saa3^−/−^* Trem2^hi^ Mφ may possess enhanced lipid uptake and metabolism capabilities, which could contribute to the significant reduction in plaque size. Additionally, genes like *Ctsk*, *Gpnmb*, *Igf1*, and *Spp1* are associated with tissue remodeling pathways, indicating a potential shift of *Saa3^−/−^*Trem2^hi^ Mφ towards an anti-inflammatory phenotype (Fig. 5b). Differential gene expression analysis utilizing violin plots demonstrated that genes associated with tissue repair and lipid metabolism—specifically *Gpnmb*, *Fabp5*, *Lpl, Lgals3*, *Cd36*, *Spp1*, and *Pparg*—are more prominently expressed in *Saa3^−/−^* Trem2^hi^ Mφ compared to their WT counterparts (Fig. 5c). CD36 is a scavenger receptor widely expressed on the surface of various cell types, including macrophages. CD36 binds to oxLDL particles, facilitating their uptake into the macrophage, particularly in the context of atherosclerosis and inflammation [98]. Concomitantly, there is a downregulation of genes in *Saa3^−/−^*than WT Trem2^hi^ Mφ such as *Thbs1, Kdm7a, Vegfa, Slc7a11, Fn1*, and *Chil3*. These genes correspond to GO enrichment pathways such as positive regulation of chemotaxis, positive regulation of vasculature development, and angiogenesis (Fig. 5b). Angiogenesis is associated with inflammatory cells and can contribute to plaque instability [99, 100]. This complex gene expression network underscores the multifaceted influence of *Saa3* in atherosclerosis and pinpoints interconnected genes geared toward modulating macrophage function to combat disease progression.

**Fig. 5.**
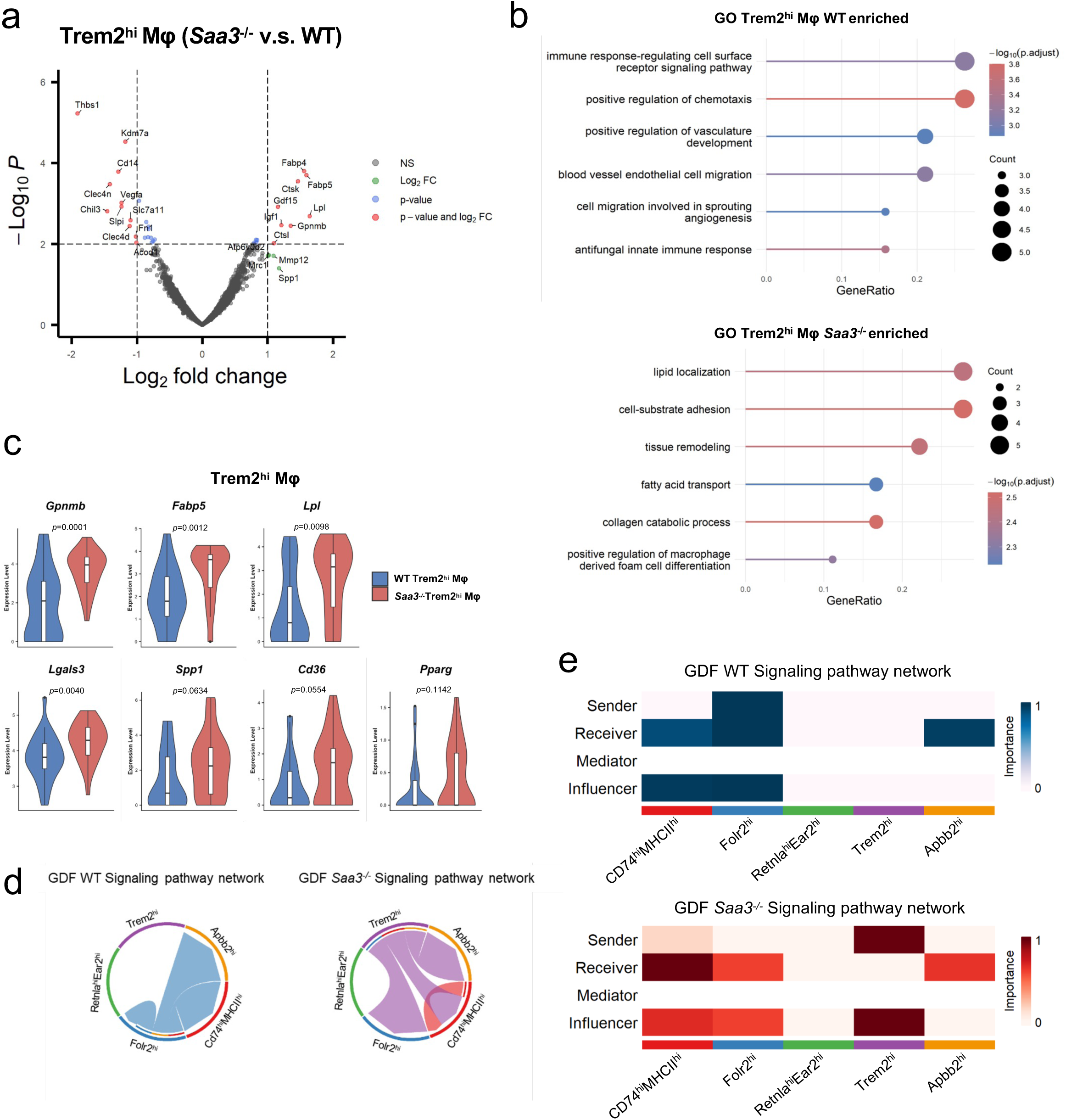
Aortic Trem2^hi^ macrophages in *Saa3^−/−^* mice express more lipid metabolism and tissue repair-associated genes. **(a)** Volcano plot illustrating differential gene expression in aortic Trem2^hi^ macrophage (Mφ) population of *Saa3^−/−^* versus WT mice. **(b)** The bubble plot presents the GO enrichment analysis for Trem2^hi^ Mφ from WT versus *Saa3^−/−^* (upper panel) or *Saa3^−/−^* versus WT (lower panel) mice. **(c)** Violin plots depicting the expression levels of tissue repair-associated genes in Trem2^hi^ Mφ from WT and *Saa3^−/−^* mice aortas. Each plot indicates the gene expression level for *Gpnmb*, *Fabp5*, *Lpl*, *Lgals3*, *Spp1*, *Cd36*, and *Pparg*, with blue representing WT and red for *Saa3^−/−^* group. The central white bar denotes the median expression level, while the thick bar within the violins indicates the interquartile range. The width of the violins reflects the density of data points at different expression levels. **(d)** The circular diagram illustrates the GDF signaling pathway networks in aortic macrophage subpopulations from WT (left) and *Saa3^−/−^* (right) mice. Each segment of the circle represents a different macrophage subpopulation. The connections between segments indicate interactions and signaling pathways shared among these subpopulations. **(e)** Heatmap representation of the inferred GDF signaling pathway network across five sub-populations of aortic macrophages. Each column represents a specific macrophage sub-population, while each row corresponds to the role of the population in the signaling pathway as a sender, receiver, mediator, or influencer.

Analysis of aortic Trem2^hi^ macrophages from WT and *Saa3^−/−^* mice also revealed a consistent expression pattern for a cohort of genes associated with lipid metabolism and cholesterol processing (Figs 5b-5c). The violin plots compare the expression levels of genes central to cholesterol efflux pathways, including *Abca1, Abcg1, Nr1h2, Scarb, Nr1h3*, and *Apoe* (Fig. 5c, Supplementary Fig. 4). There were no statistically significant differences observed between the two genotypes for most genes (Fig. 5c).

Analysis through *CellChat* has additionally revealed alterations in the growth differentiation factor (GDF) signaling pathway in all subsets of the aortic macrophages (Figs. 5d-5e). In the *Saa3^−/−^* Trem2^hi^ macrophage phenotype, there is a notable shift towards the sender role in the GDF signaling network (Fig. 5d). In contrast, Folr2^hi^ macrophages predominantly assume the signaling initiative in the WT condition (Figs. 5d-5e). This shift in signaling behavior suggests that the absence of *Saa3* leads to a reprogramming of the Trem2^hi^ macrophage population, endowing them with a more prominent role in initiating signaling within the GDF pathway.

GDF-15 has been reported to reduce cardiac damage and rupture after myocardial infarction by mitigating inflammation and apoptosis via the PI3K/Akt pathway. It also protects against atherosclerosis by modulating apoptosis, inhibiting monocyte recruitment and macrophage activation, and enhancing plaque stability [101–103].

Other studies suggest GDF-15 may reduce macrophage accumulation in *Ldlr^−/−^* or *Apoe^−/−^* mice [104–106]. This shift in the gene expression landscape highlights the critical role of *Saa3* in regulating macrophage function and may provide insight into the molecular mechanisms underpinning the immune responses in aortic plaque progression.

### SAA3 modulates cholesterol efflux, synthesis, and inflammatory responses in macrophages under steady-state condition or during foamy macrophage differentiation

SAA3 has been shown to exacerbate atherosclerosis in *ApoE^−/−^* mice, as evidenced by increased lesion area upon overexpression and reduced disease severity with suppression [20]. Another study also observed reduced aortic plaque lesions in atherosclerotic *Saa3^−/−^Ldlr^−/−^* mice and indicated that these effects are sex-dependent [14]. To validate our scRNA-seq observations on aortic macrophages and investigate the role of *Saa3* in foamy macrophages, we performed qRT-PCR to assess the expression of inflammatory genes (*Il1b* and *Tnf*), cholesterol efflux genes (*Abca1* and *Abcg1*), and cholesterol synthesis genes (*Cyp51* and *Hmgcr*) in bone marrow-derived macrophages (BMDMs) from WT and *Saa3^−/−^* mice (Fig. 6a). These BMDMs were treated overnight with either media alone or 20 µg/ml soluble cholesterol to differentiate cells into foamy macrophages (Fig. 6a). We also conducted scRNA-seq analysis on WT BMDMs treated with SAA3 in either media alone or 20 µg/ml soluble cholesterol (Fig. 6a). Overall, gender-specific effects were generally minimal under these experimental conditions, while genes involved in cholesterol efflux and synthesis were significantly lower in female WT foamy macrophages treated with SAA3 compared to males (Figs. 6b-c). Consistent with the previous study [14], *Il1b* expression was significantly upregulated in BMDMs from female *Saa3^−/−^* mice compared to WT under media-alone conditions, with no such effect observed in male mice, while *Tnf* expression remained unchanged between genotypes (Fig. 6b).

**Fig. 6.**
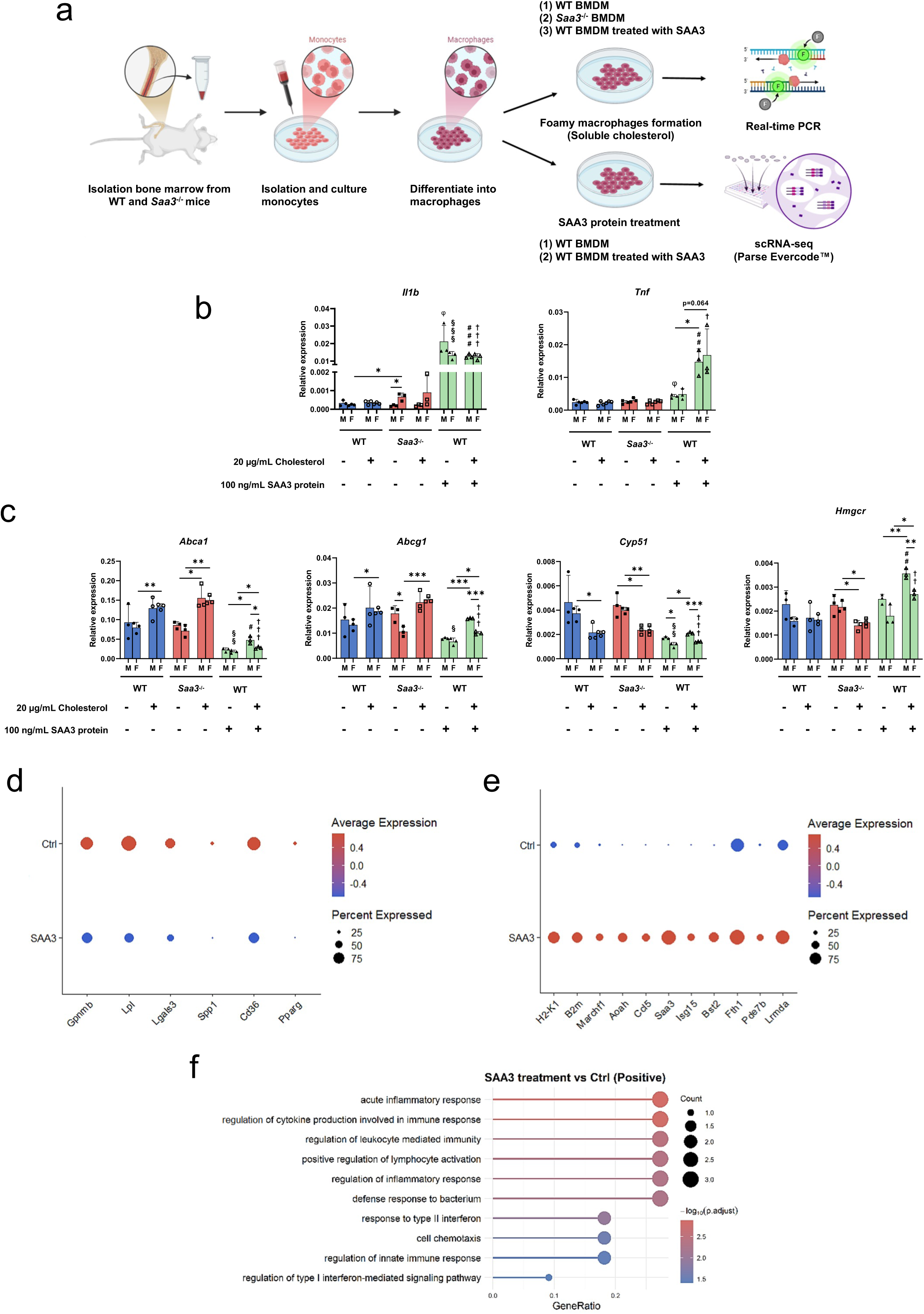
SAA3 modulates cholesterol metabolism, inflammatory gene expression, and immune response pathways in macrophages under steady-state condition or during foamy macrophage differentiation. **(a)** Schematic overview of the experimental design. Bone marrow cells were isolated from WT and *Saa3^−/−^*mice. Then cultured and differentiated into bone marrow-derived macrophages (BMDMs), followed by treatment with soluble cholesterol to induce foamy macrophages or recombinant SAA3 protein. Downstream analyses included real-time PCR and single-cell RNA sequencing (scRNA-seq) using Parse Evercode™. **(b-c)** As treatment conditions in **(a)**, Real-time PCR analysis of genes involved in inflammatory cytokines (*Il1b*, *Tnf*) and cholesterol metabolism (*Abca1*, *Abcg1*, *Cyp51*, *Hmgcr*) and under various treatment conditions. Data are represented as relative gene expression normalized to the control group, with significant differences marked by asterisks (\**p* < 0.05, \*\**p* < 0.01, \*\*\**p* < 0.001). The symbols represent the following comparisons: **φ** indicates SAA3-treated male WT BMDMs versus male WT BMDMs cultured in media alone, **§** indicates SAA3-treated female WT BMDMs versus female WT BMDMs cultured in media alone, **#** indicates SAA3-treated male WT BMDMs in the presence of soluble cholesterol versus male WT BMDMs in soluble cholesterol media alone, and **†** indicates SAA3-treated female WT BMDMs in the presence of soluble cholesterol versus female WT BMDMs in soluble cholesterol media alone. The number of symbols denotes the level of statistical significance, where one symbol indicates *p* < 0.05, two symbols indicate *p* < 0.01, and three symbols indicate *p* < 0.001. **(d-f)** scRNA-seq analysis of WT BMDMs treated with media alone or SAA3 protein. **(d-e)** Dot plots show average expression (color intensity) and the percentage of cells expressing key genes (dot size) related to tissue repair, lipid metabolism, and inflammatory pathways. **(f)** Gene Ontology (GO) enrichment analysis of pathways upregulated in SAA3-treated BMDMs compared to controls. The size of the dots represents the gene ratio, and color intensity represents statistical significance (-log_10_ (adjusted *p*-value)). Statistical significance was determined using ANOVA or t-tests where appropriate.

Following soluble cholesterol treatment, both WT and *Saa3^−/−^* BMDMs present minimal differences, with upregulated expression of the cholesterol efflux gene (*Abca1*) and downregulated expression of the cholesterol synthesis gene (*Cyp51*), indicative of the foamy macrophage phenotype (Fig. 6c). SAA3 treatment significantly reduced cholesterol efflux gene expression (*Abca1* and *Abcg1*) while increasing cholesterol synthesis gene expression (*Hmgcr*) in WT BMDMs undergoing foamy macrophage differentiation (Fig. 6c). We next performed scRNA-seq analysis on SAA3-treated WT BMDMs to confirm that genes associated with tissue repair and lipid metabolism (*Gpnmb*, *Lpl*, *Lgals3*, *Spp1*, *Cd36*, *Pparg*) were significantly downregulated compared to untreated controls (Fig. 6d). This aligns with the *in vivo* phenotypes, where Trem2^hi^ aortic macrophages from *Saa3^−/−^* mice exhibited significantly higher expression of these genes compared to Trem2^hi^ macrophages from WT mice (Figs. 5c, 6d). We further observed the genes associated with acute inflammatory response, interferon responses, and cell chemotaxis are increased in SAA3-treated WT BMDMs (Figs. 6e-6f). Notably, exogenous SAA3 significantly increases *Il1b* expression in both foamy and non-foamy macrophages (Fig. 6b).

These data indicate that SAA3 modulates inflammatory and lipid metabolism pathways in macrophages, particularly during foamy macrophage differentiation, by reducing cholesterol efflux, increasing cholesterol synthesis, and enhancing inflammatory and interferon responses, suggesting that SAA3 potentially exacerbates atherosclerosis progression through these mechanisms.

## Discussion

Gut microbiota plays significant roles in inducing and maintaining sex dimorphism, particularly in relation to immune responses, metabolism, and hormone regulation. In NOD mice, the sex difference in type 1 diabetes susceptibility disappears under germ-free conditions, while gut microbiota transfer from males to females elevates testosterone levels and protects against disease [107]. This protection, mediated by microbiota-regulated sex hormones, is lost upon androgen receptor blockade[107].

Metabolic syndrome is also mediated by the gut microbiota in a sex-dependent manner, as demonstrated by the transfer of this sex-specific effect through fecal transplantation between male and female mice, with antibiotic treatment eliminating these differences and highlighting the causative role of microbiota in sexual dimorphism [108]. Furthermore, the microbiota regulates bone marrow mesenchymal stem cells (BMMSCs) by inhibiting osteogenesis and promoting adipogenesis, as shown by increased osteogenic differentiation in germ-free (GF) mice, while colonization with specific-pathogen-free (SPF) microbiota restored normal BMMSC proliferation, differentiation, and immunomodulatory functions [109]. The microbiota regulates hematopoietic stem cell (HSC) self-renewal and differentiation under stress by modulating local bone marrow iron availability through red blood cell recycling by macrophages [110]. Notably, the gut microbiota modulates the expression of SAA3 in colonic and adipose tissue via Toll-like receptor signaling [15]. These findings suggest that the gut microbiota may influence confounding factors, such as iron availability and SAA3 levels, within the tissue microenvironment, thereby affecting sex dimorphism and immune cell differentiation. The SAA3-induced sex dimorphism observed in previous studies may also be influenced by microbiota-driven effects on bone marrow cell differentiation during early life.

Previous study indicates that SAA3 promotes atherosclerosis in male mice while providing protective effects in female mice [14]. Jylhava et. al. suggest a potential sex-dimorphism in the relationship between serum amyloid A (SAA) and early atherosclerosis, as SAA levels correlated with carotid intima-media thickness (IMT) in men but not in women, indicating differing underlying mechanisms of atherosclerotic risk between the sexes [111]. The absence of SAA3 in female mice led to a significant decrease in adipose tissue macrophage content and inflammation, while similar effects were not observed in male mice [112]. Another study highlights a significant sex dimorphism in the association between SAA levels and adipocyte size, with pronounced correlations observed in lean and overweight women, while no such associations were found in men [113]. Our results indicate there is still less aortic plaque progression in female atherosclerotic *Saa3^−/−^* than in WT mice (Fig. 1).

The gender differences were minimal but SAA3 significantly reduced cholesterol efflux genes and increased synthesis genes, particularly in female WT foamy macrophages (Fig. 6). SAA3-treated WT BMDMs showed downregulation of tissue repair and lipid metabolism genes and upregulation of acute inflammatory and interferon-response genes, aligning with *in vivo* phenotypes of Trem2^hi^ macrophages in *Saa3^−/−^* mice represented the higher expression of lipid metabolism genes (Fig. 6). These results suggest that SAA3 modulates inflammatory and lipid pathways in foamy macrophages, which may be exacerbating atherosclerosis progression *in vivo*. The single-cell RNA sequencing (scRNA-seq) analysis of CD45+ cells from aortic arches revealed significant alterations in the aortic immune cell landscape of *Saa3^−/−^*mice. Increased proportions of T cells, B cells, monocytes, and γδ T cells were noted, while macrophages and innate lymphoid cells (ILCs) were reduced compared to WT mice (Fig. 2). This shift in immune cell composition suggests that *Saa3* deficiency alters the inflammatory responses within the aortic arches, potentially contributing to the observed reduction in plaque progression. *CellChat* analysis further highlighted the impact of *Saa3* deficiency on intercellular communication within the aortic microenvironment (Figs. 2, 3). *Saa3^−/−^* mice exhibited enhanced intercellular interaction strength, particularly among γδ T cells and all other immune cell types, suggesting a more reactive immune milieu. Conversely, the reduced macrophage-to-all other immune cell signalings were observed, indicating a dampened inflammatory response (Fig. 2). These findings suggest that *Saa3* deficiency promotes shifts towards reparative and less inflammatory phenotypes in aortic immune cells, potentially influencing atherosclerosis progression.

Compared to our previously published dataset, where we profiled single-cell sorted *Cx3cr1*-derived aortic macrophages during atherosclerosis progression and regression [8], we identified 11 aortic macrophage clusters, while we only identified five distinct clusters with notable differences in gene expression between WT and *Saa3^−/−^*mice (Fig. 3a). This may be due to the difference in the platform used to capture single cells between 10X Genomics and BD Rhapsody. Nonetheless, we still observed that Trem2^hi^ macrophages were more abundant in *Saa3^−/−^* mice (Fig. 3b) and revealed gene expression profiles indicative of enhanced lipid metabolism and tissue repair functions (Fig. 5). These macrophages may be crucial in mitigating plaque progression by promoting an anti-inflammatory environment and enhancing extracellular matrix production. Our findings also highlight the differential ligand-receptor interactions in WT and *Saa3^−/−^* mice macrophages (Supplementary Fig. 3). *Saa3* deficiency led to altered signaling dynamics, particularly in Trem2^hi^ macrophages, which showed increased interactions involving *Spp1* and integrins, as well as type IV collagen, suggesting enhanced adhesion and migration capabilities. In contrast, chemokine-mediated communications were more prominent in WT macrophages, facilitating the recruitment of inflammatory cells to atherosclerotic lesions (Supplementary Fig. 3d). The observed changes in gene expression and intercellular communication underscore the complex role of *Saa3* in modulating Trem2^hi^ macrophage functions during atherosclerosis. *Saa3* deficiency appears to shift Trem2^hi^ macrophages towards a phenotype that promotes tissue repair and plaque stability (Fig. 5), which may explain the reduced plaque progression despite elevated cholesterol levels.

In conclusion, our study reveals that *Saa3* deficiency has a dual impact on atherosclerosis, reducing aortic plaque progression in female mice while increasing peripheral lipid levels. The enhanced reparative and anti-inflammatory properties of macrophages in *Saa3^−/−^* mice likely contribute to the observed protective effects against atherosclerosis. *Saa3* is exclusively expressed in macrophages within the immune cell population and SAA3 modulates inflammatory and lipid pathways in foamy macrophages. These findings provide valuable insights into the regulatory mechanisms of immune function within the aortic niche and highlight the potential of targeting *Saa3* for therapeutic strategies in atherosclerosis. Further research is warranted to elucidate the precise molecular pathways involved and to explore the therapeutic potential of modulating *Saa3* expression in cardiovascular diseases. The observed alterations in immune cell composition and intercellular communication in *Saa3*-deficient mice highlight a shift toward reparative and anti-inflammatory phenotypes, potentially stabilizing plaques despite elevated cholesterol levels.

### Author contributions

TYC, YZY, and JDL designed experiments and interpreted the data. TYC and YZY wrote the manuscript draft, followed by JDL editing. TYC, YZY, CJL, JCW, CHY, YTC, and PAC performed experiments. TYC and YZY coordinated and performed the single-cell RNA-seq experiments and then analyzed single-cell RNA-seq data.

## Acknowledgments

We thank the Animal Resource Center and the Consortium of Integrative Biomedical Science Key Technology at National Taiwan University for their technical support. We also acknowledged the AAV Core Facility of Academia Sinica in generating recombinant AAV (Grant AS-CFII112-204) and National Center for High-performance Computing (NCHC) of National Applied Research Laboratories (NARLabs) in Taiwan for providing computational and storage resources. Funding: This work was financially supported by the National Science and Technology Council, Taiwan (110-2636-B-002-021, 111-2636-B-002-024, 112-2636-B-002-015, 113-2636-B-002-009), National Taiwan University (113L894501, 113-UN0066), Ministry of Education (Yushan Young Fellow Program, Featured Areas Research Center Program of “Center for Advanced Computing and Imaging in Biomedicine” (NTU-113L900701))

## Material and Methods

### Animals

All animal procedures were approved by the National Taiwan University IACUC Committee. The Saa3-knockout mice (*Saa3^−/−^*; C57BL/6N-Saa3tm1.1(KOMP)Vlcg/MbpMmucd) were developed by the Mutant Mouse Resource & Research Centers (MMRRC). The ES cell clone 13807A-A1 was injected into morulae or blastocysts to generate chimeric mice. These chimeras were then bred with C57BL/6N mice, resulting in the production of heterozygous tm1 (Deletion) mice. To achieve recombination at the LoxP sites, heterozygous tm1 mice were crossed with a ubiquitous Cre deleter mouse line, generating tm1.1 (CRE-mediated Deletion) allele mice. Notably, the archived germplasm does not carry the Cre transgene. Cryo-recovery of the strain will be conducted using C57BL/6N females by the MMRRC. Detailed information about the Cre-mediated deletion process can be found at the following reference: Velocigene KOMP Repository, clone 13807, http://velocigene.com/komp/detail/13807. The cryopreserved sperm were acquired and recovered by the National Laboratory Animal Center in Taiwan. We generated heterozygous *Saa3^+/-^* mice and bred them to obtain 12-week-old Wild-type C57BL/6J mice wild-type (WT) and *Saa3^−/−^* littermates used for the experiments. Murine atherosclerosis was induced by injecting intraperitoneally once with AAVmPCSK9 (ssAAV8/hAAT-mPCSK9D377Y, AAV core, Institute of Biomedical Sciences, Academia Sinica, using a plasmid acquired from Addgene (Plasmid #58376) and kindly provided by Dr. Mi-Hua Tao) at 5×10^11^ viral particles/mouse and fed on Western diet (Research Diets, Cat# D12079Bi) for 18 weeks. All mice were housed in a specific pathogen-free (SPF) environment within individually ventilated cages (IVC) at a ventilation rate of 70-100 air changes per hour (ACH) under positive pressure. The facility maintained a constant temperature of 22 ± 2°C and a relative humidity of 55 ± 10%. A 12-hour light-dark cycle was regulated by timers (7:00 AM to 7:00 PM light, 7:00 PM to 7:00 AM dark).

### Cholesterol level measurements

Blood samples were collected from mice and total cholesterol levels in plasma were assessed in WT and *Saa3^−/−^* mice subjected to AAVmPCSK9D377Y and fed on Western Diet for 0, 4, 8, 12, and 16 weeks (n=5 mice per group). Cholesterol levels were measured using a colorimetric assay (FUJIFILM/Wako, Cat# 639-50981), with absorbance read at 600 nm following a 5-minute incubation at 37 °C. HDL and LDL/VLDL cholesterol levels in plasma were quantified using the ab65390 Cholesterol Assay Kit (Abcam, Version 13a, Updated 17 April 2023), following the manufacturer’s protocol with adaptations for our setup. Plasma (20-30 μL) was mixed with an equal volume of 2X Precipitation Buffer to separate the HDL and LDL/VLDL cholesterol fractions. After adding reaction mixes, samples were incubated at 37 °C for 60 minutes away from light. Absorbance was measured at 570 nm using a microplate reader, and cholesterol concentrations were determined by correlating absorbance readings with the standard curve. Statistically significant differences between the two groups at respective time points were determined using the unpaired Mann-Whitney U-test (*p < 0.05).

### Histology

At harvesting, the aortic arch was meticulously excised from the experimental mice for the subsequent single-cell isolation, and then aortic roots were embedded in OCT (Catalog # NC0696746). Serial sections were performed at 6 μm on embedded aortic roots and stained with Oil Red O solution (Sigma) for 10 minutes at 37°C. Intimal lesions and stained areas were imaged by an inverted fluorescence microscope (OLYMPUS CKX53) and quantified by ImageJ ^17^.

### Aortic cells flow cytometry sorting and single-cell RNA-seq library preparation

The protocol for isolating aortic cells from mice is as previously described^8^. Briefly, the procedure began with aorta perfusion using phosphate-buffered saline (PBS). After perfusion, the aortas were stored in FACS buffer (1X HBSS, 1% BSA, 1mM EDTA, 20mM HEPES, and 1mM Sodium Pyruvate). The enzymatic digestion processes involved immersing the aorta in a digestion buffer containing 4U/ml liberase (Roche, Cat. 54001127001), 100 ug/ml hyaluronidase (Sigma, Cat. H3506), and 100 ug/ml DNase I (Sigma, Cat. DN25), followed by fine mincing. The mixture was incubated at 37°C to facilitate effective digestion, with the process repeated 5-6 times to ensure complete tissue breakdown. After digestion, the mixture was filtered through a 35µm strainer to obtain a single-cell suspension. The collected cells were then stained with CD45 (Alexa Fluor 488 anti-mouse CD45, Clone: 30-F11, Biolegend, Cat. 103122) and Live/Dead (L/D) Blue markers (Thermo Fisher, Cat. L34962), and the CD45-positive, L/D Blue-negative cells were sorted as Supplementary Figure 1 to prepare single-cell libraries.

Single-cell samples were labeled using the BD™ Mouse Immune Single-Cell Multiplexing Kit (Cat. 633793) for each cartridge run on the BD Rhapsody Express Single Cell Analysis System for single-cell capture, followed by cDNA synthesis with the BD Rhapsody cDNA Kit (Cat. 633773). The mRNA Whole Transcriptome Analysis (WTA) and Sample Tag libraries were prepared using the BD Rhapsody™ WTA Amplification Kit (Cat. 633801), according to the manufacturer’s guidelines. The quality and quantity of the libraries were assessed using the Agilent 4200 TapeStation System with Agilent High Sensitivity D1000 ScreenTape (Agilent Technologies, Santa Clara, CA) and a Qubit Fluorometer with the Qubit dsDNA HS Assay (Invitrogen, MA, USA). Finally, the libraries were sequenced using the NovaSeq PE150 sequencer (Illumina, San Diego, CA).

### Aortic single-cell sequencing data reads alignment and analysis

The fastq files generated from sequencing were processed using Seven Bridges Genomics. The data were aligned to the mouse_GRCm38-PhiX-gencodevM19-20181206 and demultiplexed using hashtags to separate samples. Subsequent steps included filtering low-quality cells and identifying unique molecule identifiers (UMIs) to produce filtered matrices, which were then utilized as inputs for further analysis. The filtered matrices were analyzed using the Seurat package (v.4.3.0)^103^ in R (v.4.3.1). Initially, to exclude low-quality cells, we filtered out cells with nFeature_RNA < 200 and percent.mt > 15, tailoring filter parameters for each sample based on violin plots of RNA feature numbers and the percentage of mitochondria in each cell. We used the FindVariableFeatures() function in each sample to select the top 2000 genes with the highest variance using the variance variance-stabilizing transformation (VST) method for downstream analysis. Next, we applied a linear transformation to scale the data, ensuring equal weight to each gene expression level in downstream analysis. A linear dimensional reduction (PCA) was performed, and the Elbow plot was used to decide which PCs to include in the following steps. We cluster cells using the Louvain algorithm (resolution = 0.5). The non-linear dimensional reduction method was employed for visualizing cell clustering, such as UMAP.

To analyze two or more single-cell datasets together, we followed the Seurat integration analysis pipeline. We used the FindIntegrationAnchors() function to identify anchors, followed by the IntegrateData() function to use these anchors to integrate the datasets. To automatically annotate cell types, SingleR (v.2.8.0) was utilized with the ImmGen database, which specifically focuses on immunological cell types. To ensure the analysis targeted the immune population, contaminating cell types such as fibroblasts and stromal cells were subsequently excluded. To identify differentially expressed features, we employed the FindAllarkers() or FindMarkers() functions to identify biomarkers for each cell cluster (min.pct = 0.25, logfc.threshold = 0.25). We used visualization tools such as VlnPlot, feature plot, dotplot, and heatmap to display intergroup gene expression differences. Additionally, to comprehend the potential biological pathway connections of gene sets expressed by each cell population and their differences between two genotypes, we conducted a Gene Ontology (GO) analysis using the ClusterProfiler package (v4.8.3)^104^. This was done after identifying differentially expressed genes (DEGs) in each cell population compared to the cell population of another genotype. The results were visualized using a Dotplot. Moreover, the analysis of intercellular communications was conducted using the CellChat package (v1.6.1)^105^. This approach allowed us to understand the roles played by each cell group in signal transduction pathways and compare the differences in cellular signaling among cell groups under different genotypes.

### Bone marrow-derived macrophages (BMDMs) culture

Bone marrow cells harvested from C57BL/6J mice were cultured in RPMI 1640 medium supplemented with 10% heat-inactivated fetal bovine serum (FBS), 1% penicillin/streptomycin/amphotericin B (P/S/A), and 20 ng/mL macrophage colony-stimulating factor (M-CSF). On day 3, the medium was replaced with 10 mL fresh medium. On day 5, an additional 2 mL of medium was added. After seven days of culture, adherent macrophages were detached using Accutase and treated with 100 ng/mL SAA3 protein. The cell suspension was adjusted to a density of 5 × 10⁶ cells/mL, seeded into a 48-well plate, and cultured for 24 hours.

### Real-time RT-PCR

Total RNA was extracted from BMDMs using TRIzol™ Reagent, and cDNA was synthesized using SuperScript™ IV Reverse Transcriptase. Real-time RT-PCR was performed with TaqMan™ Fast Advanced Master Mix on a CFX Connect Real-Time PCR Detection System. The assays utilized the following TaqMan probes: *Gapdh* (Mm99999915_g1), *Saa3* (Mm00441203_m1), *Il-1β* (Mm00434228_m1), and *Tnf* (Mm00443258_m1). Additionally, real-time RT-PCR was also conducted using iTaq™ Universal SYBR® Green Supermix with primers targeting *Abca1*, *Abcg1*, *Hmgcr*, and *Cyp51*. The fold change in each target mRNA was determined by normalizing its expression level to *Gapdh* within the same sample. Relative gene expression was calculated using the comparative method (2^−ΔΔCT^).

### BMDM single-cell RNA library preparation and sequencing

BMDMs were fixed and permeabilized according to the Evercode™ Fixation v2 protocol (Parse Biosciences). The split-pool barcoding process, following the Evercode™ WT v2 protocol, consisted of three rounds. In Round 1, cells were distributed into wells, and sample-specific barcodes were applied to the fixed cells using an in-cell reverse transcription (RT) reaction. Cells from all wells were then pooled together. In Round 2, cells were redistributed across a plate, and an in-cell ligation was performed to append the second barcode. The cells from each well were again pooled together. In Round 3, cells were split across a plate, and a third barcode was appended with another in-cell ligation. After pooling cells from each well, the pooled cells were divided across multiple sublibraries. The cells were then lysed, and the library was fragmented, end-repaired, and went through A-tailing followed by the ligation of the sequencing adaptor. The library was then labeled with a fourth sublibrary-specific barcode. The quality of dsDNA libraries was analyzed using the High Sensitivity D5000 ScreenTape Kit (Agilent) and concentrations were assessed with the Qubit dsDNA HS Kit (Thermo Fisher Scientific). Libraries were sequenced on a NovaSeq X Plus (Illumina) with 10% PhiX.

### BMDM single-cell RNA data processing

The sequencing data was demultiplexed and mapped to database (mm10, GRCm38) with the commercial pipeline (ParseBiosciences-Pipeline.v1.2.1). The sequence reads from different samples was then distributed based on different Round 1 barcode with R(v4.4.2). To exclude low-quality cells, we filtered cells with nFeature_RNA > 200, nFeature_RNA < 2000, and <u>percent.mt</u> < 45, adjusting the filter parameters for each sample based on violin plots of RNA feature counts and mitochondrial percentages per cell. The FindVariableFeatures() function was used to select the top 2000 genes with the highest variance, applying the variance-stabilizing transformation (VST) method for downstream analysis. The data were then scaled using a linear transformation to ensure equal weighting of gene expression levels in subsequent analyses. Principal component analysis (PCA) was performed for linear dimensional reduction, and the Elbow plot was used to determine the number of principal components (PCs) to retain. Cell clustering was conducted using the Louvain algorithm with a resolution of 0.5. For visualization, we applied the non-linear dimensional reduction method UMAP. For integrating two or more single-cell datasets, we followed the Seurat integration analysis pipeline. The IntegrateLayers() function with the CCAIntegration method was used for dataset integration, followed by the JoinLayers() function to merge the layers. Cell types were annotated automatically using SingleR (v2.8.0) with the ImmGen database, which specializes in immunological cell types. Contaminating cell types, such as fibroblasts, were excluded to ensure the analysis focused on the immune population. To identify potential biological pathway connections and differences following SAA3 protein treatment, we performed Gene Ontology (GO) analysis using the ClusterProfiler package (v4.10.1) on differentially expressed genes (DEGs) from BMDMs under different conditions. Results were visualized using DotPlot representations.

### Data and code availability

Raw sequence data from single-cell RNA sequencing experiments are deposited in the NCBI Sequence Read Archive under BioProject accession number PRJNA1129034 and gene expression omnibus (GEO) accession number GSE270922. All processing was performed in R and analysis scripts can be found on Github at (https://github.com/JDLinLab/Saa3KO-Arch-scRNAseq-pepline-)

## Supplementary Figure Legends

**Supplementary Fig. 1.**
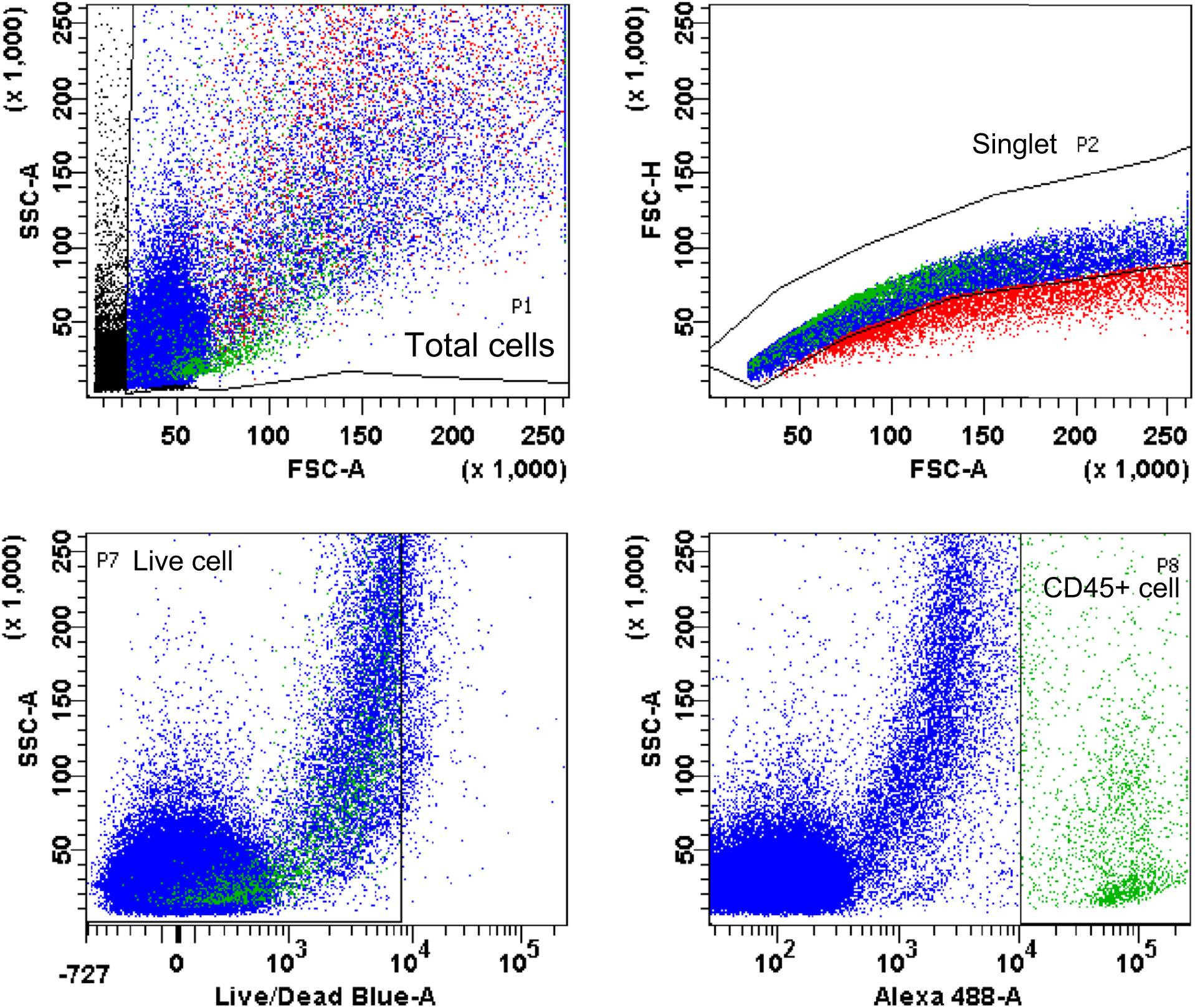
Sorting strategy of aortic CD45+ cells. Flow cytometry analysis was performed for cell sorting to isolate CD45+ immune cells from aortic arches stained for CD45 Alexa 488 and Live/Dead Blue. The top left plot shows forward scatter (FSC-A) versus side scatter (SSC-A), differentiating cells based on size and granularity, with a population gated as P1. The top right plot presents FSC-A versus FSC-H, with the discrimination gate P2 to exclude cell aggregates. The bottom left plot displays SSC-A against the Live/Dead Blue, with live cells gated as P7. The bottom right plot demonstrates SSC-A against the fluorescence intensity of CD45 Alexa-488, with the CD45+ population gated as P8. The CD45+ cells were sorted from P8 gated population.

**Supplementary Fig. 2.**
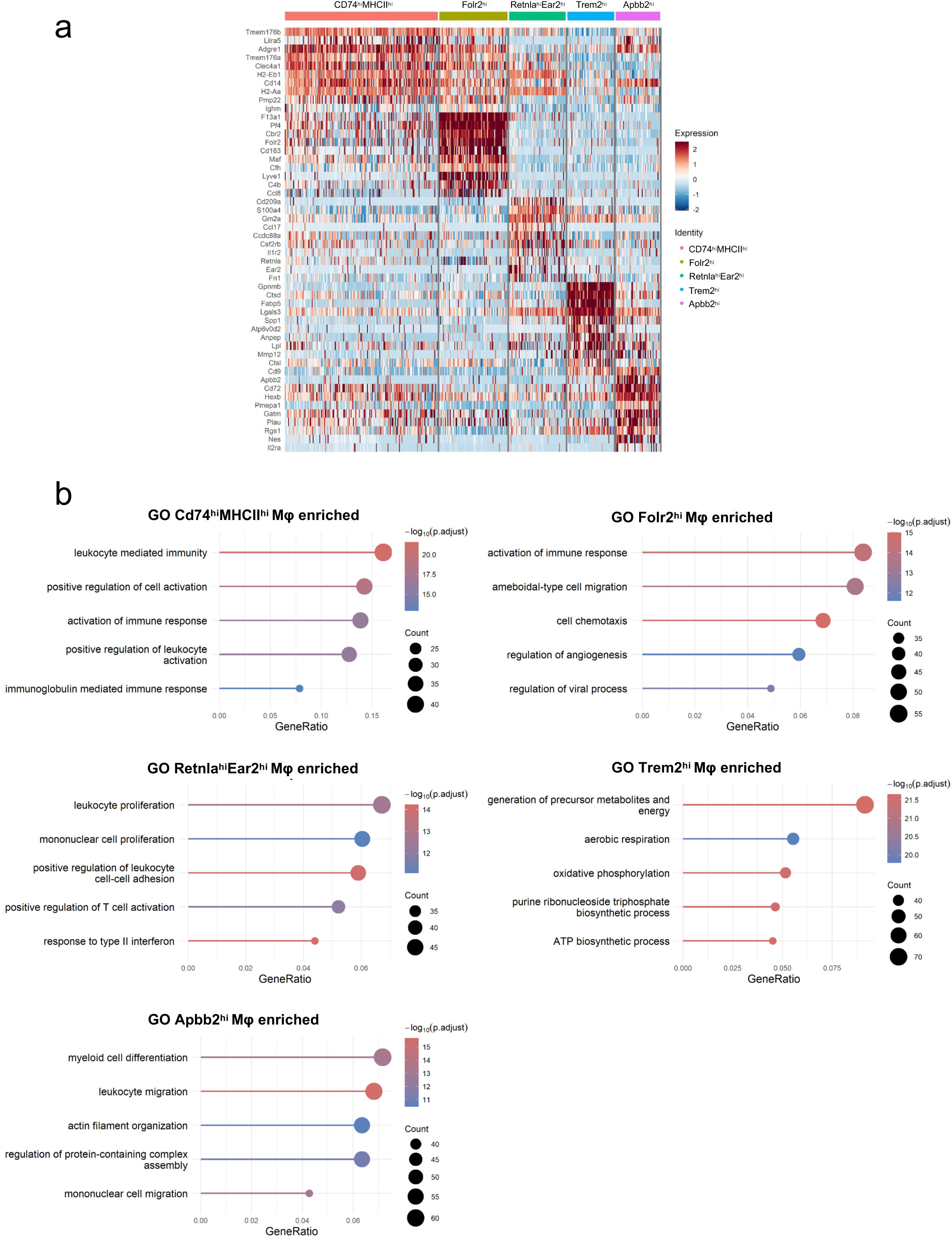
Heatmap of top 10 gene markers and GO enrichment analysis of five aortic macrophage sub-cluster. **(a)** Heatmap of top 10 marker genes of each sub-macrophage population in aortic arches from WT and *Saa3^−/−^* mice. The selection of these top marker genes was based on the results of a Wilcoxon test for differential expression. **(b)** Gene Ontology (GO) enrichment analysis of biological pathways in 5 sub-populations of aortic macrophage (Mφ), using differentially expressed genes (DEGs) identified when comparing one of specific Mφ sub-population against the rest of all other Mφ sub-populations. It can represent the functional characteristics of each Mφ sub-population.

**Supplementary Fig. 3.**
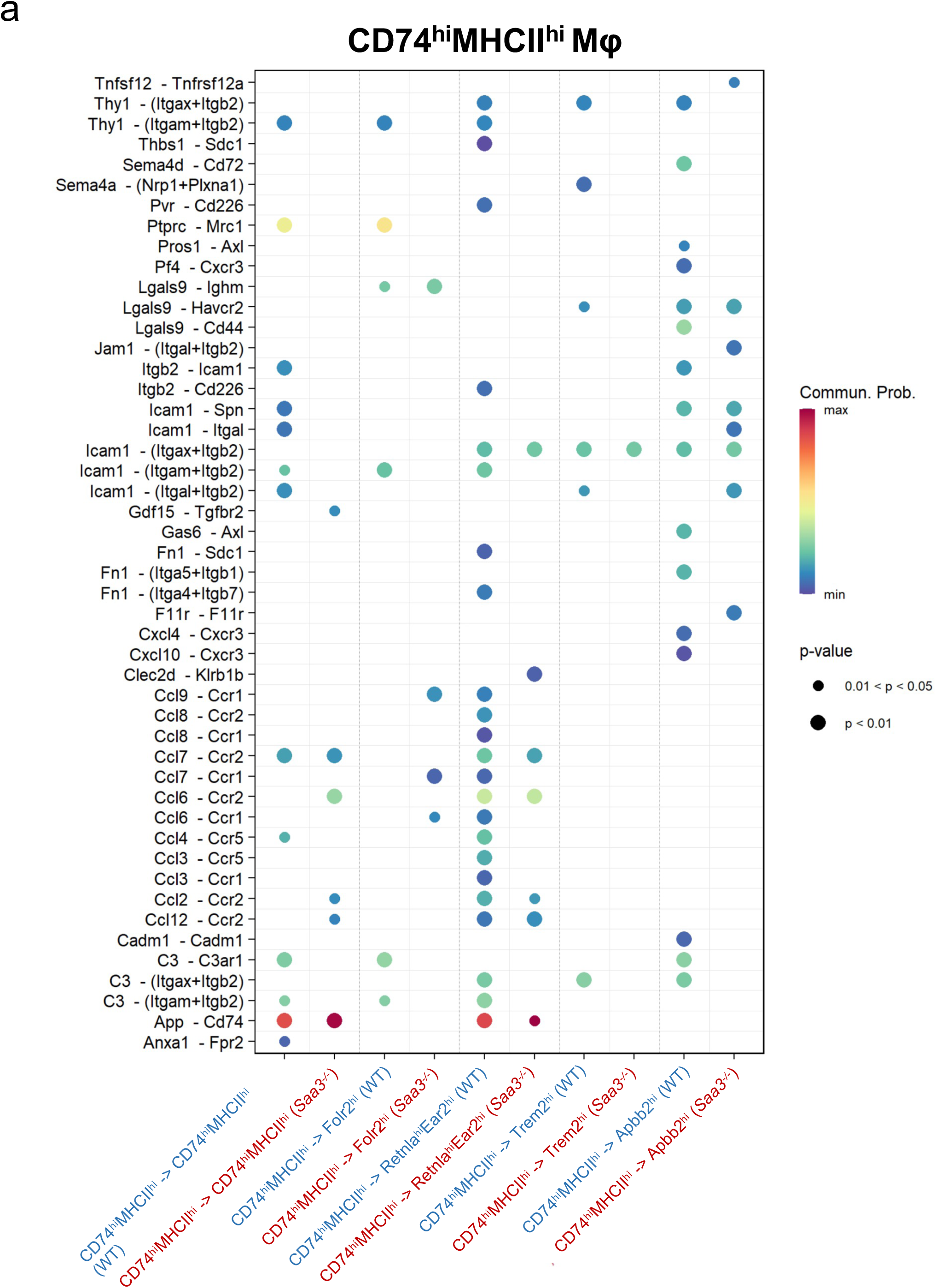
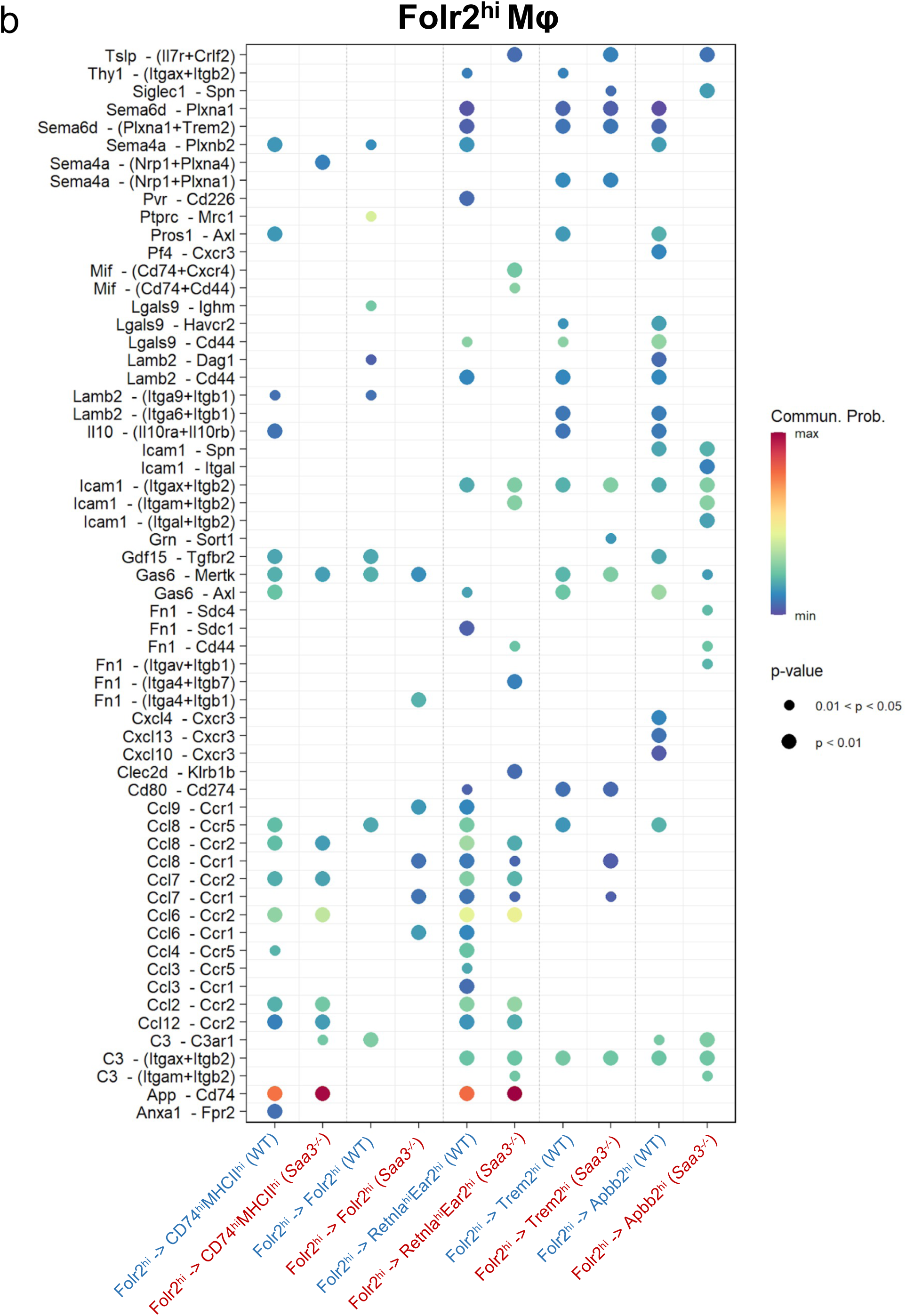
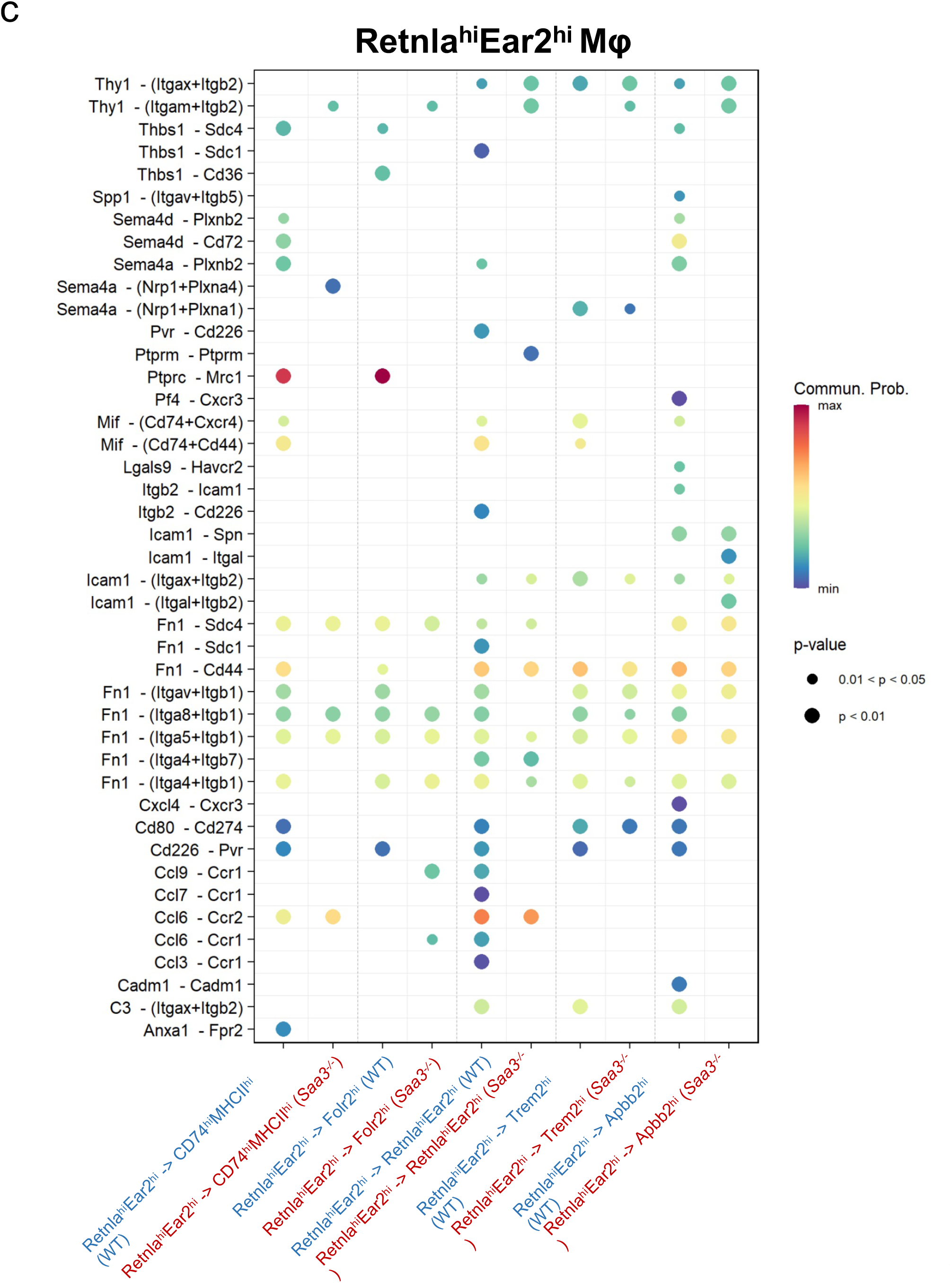
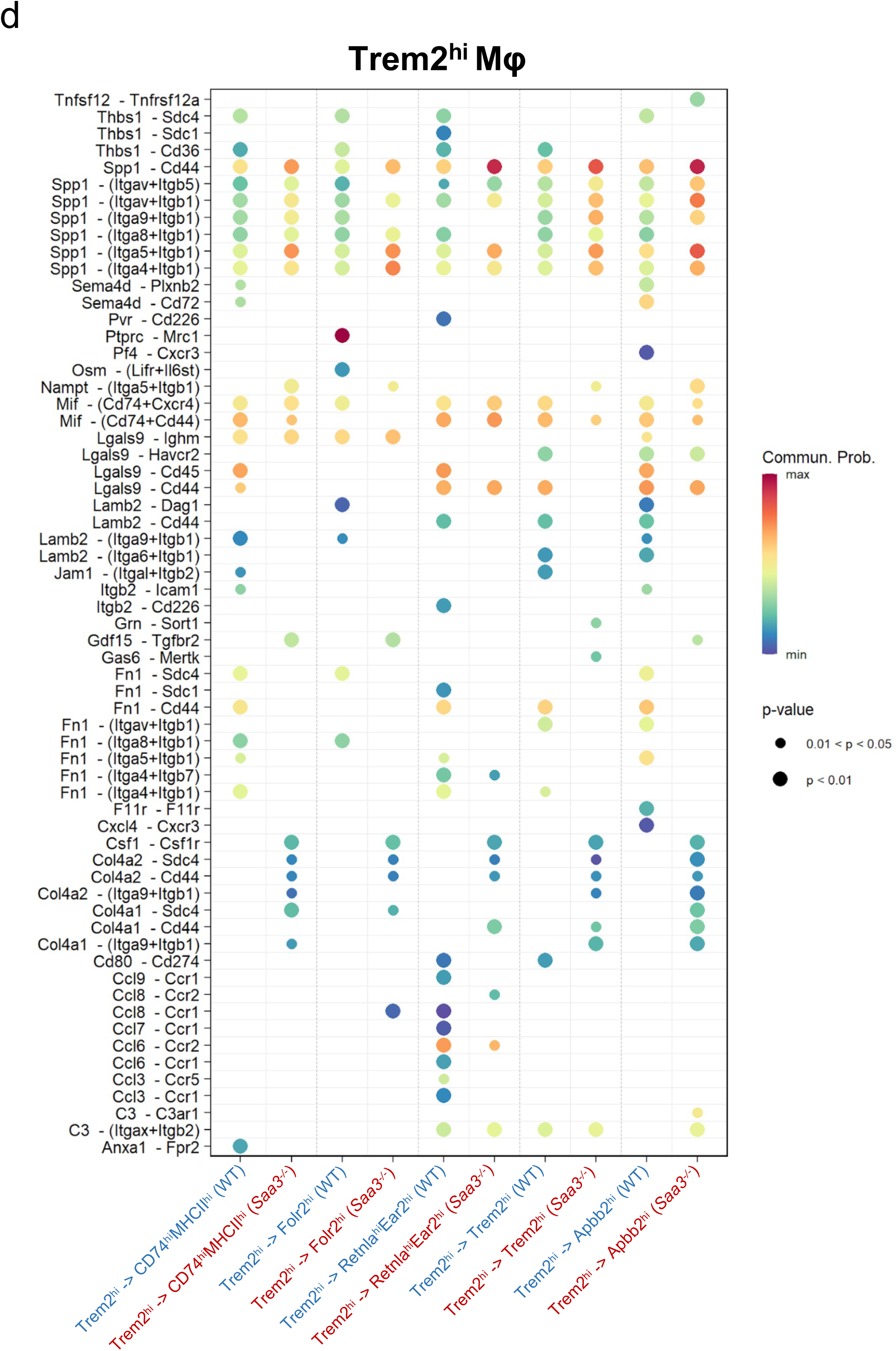
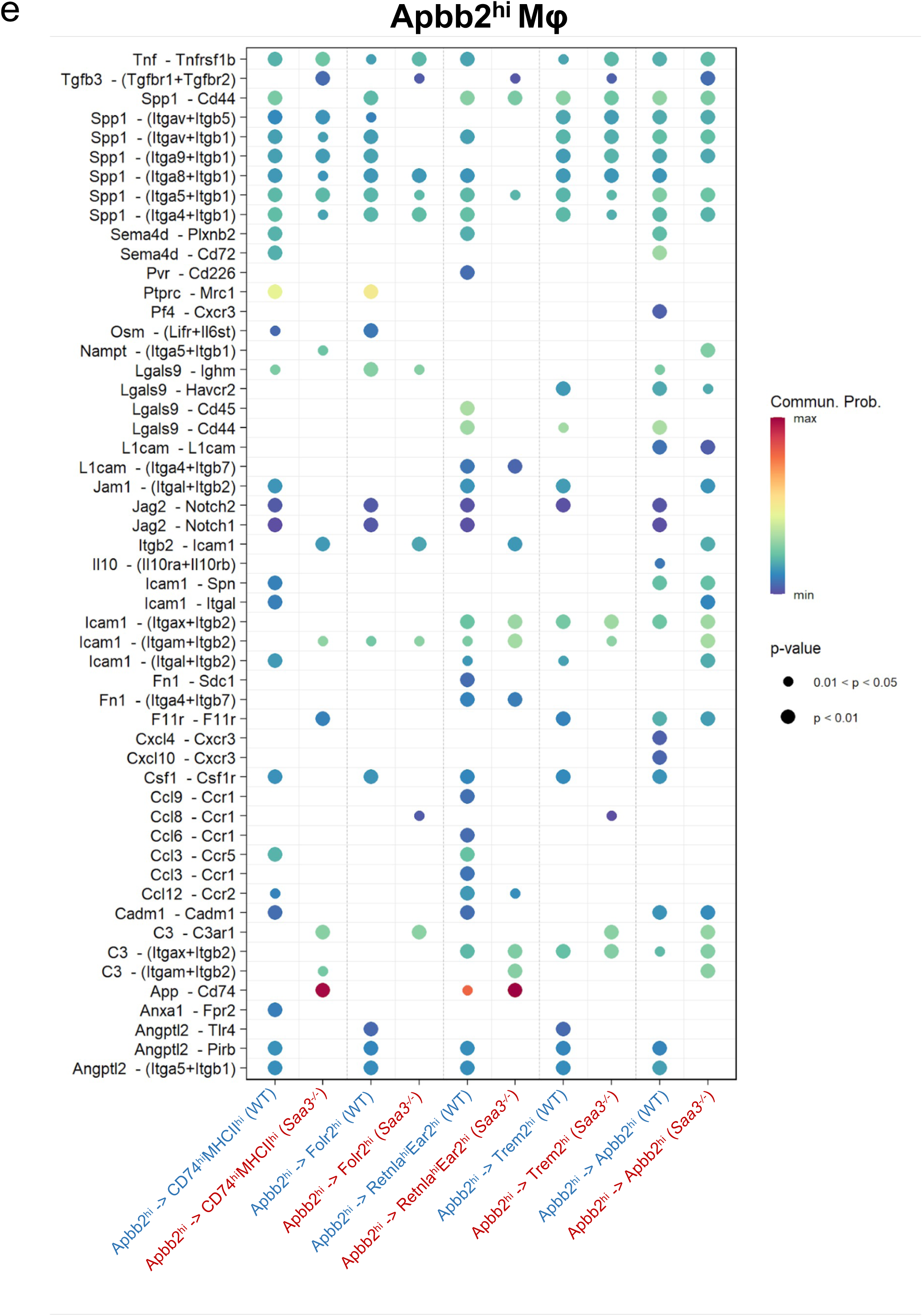
Comparison of ligand-receptor pair differences across five sub-macrophage populations between WT and *Saa3^−/−^* mice. The bubble plot illustrating the protein-protein interaction probabilities between various ligand-receptor pairs across **(a)** CD74^hi^ MHCII^hi^ macrophage (Mφ), **(b)** Folr2^hi^ Mφ, **(c)** Retnla^hi^ Ear2^hi^ Mφ, **(d)** Trem2^hi^ Mφ, and **(e)** Apbb2^hi^ Mφ between WT and *Saa3^−/−^* mice. The x-axis categorizes the conditions under which the interactions were assessed. The y-axis lists the ligand-receptor pairs. Each bubble’s color corresponds to the communication probability, with a gradient from minimum (blue) to maximum (red). The communication probability can be seen as a numerical value representing how strong or likely this interaction is between specific cell types. The size of each bubble represents the *p*-value of the interaction.

**Supplementary Fig. 4.**
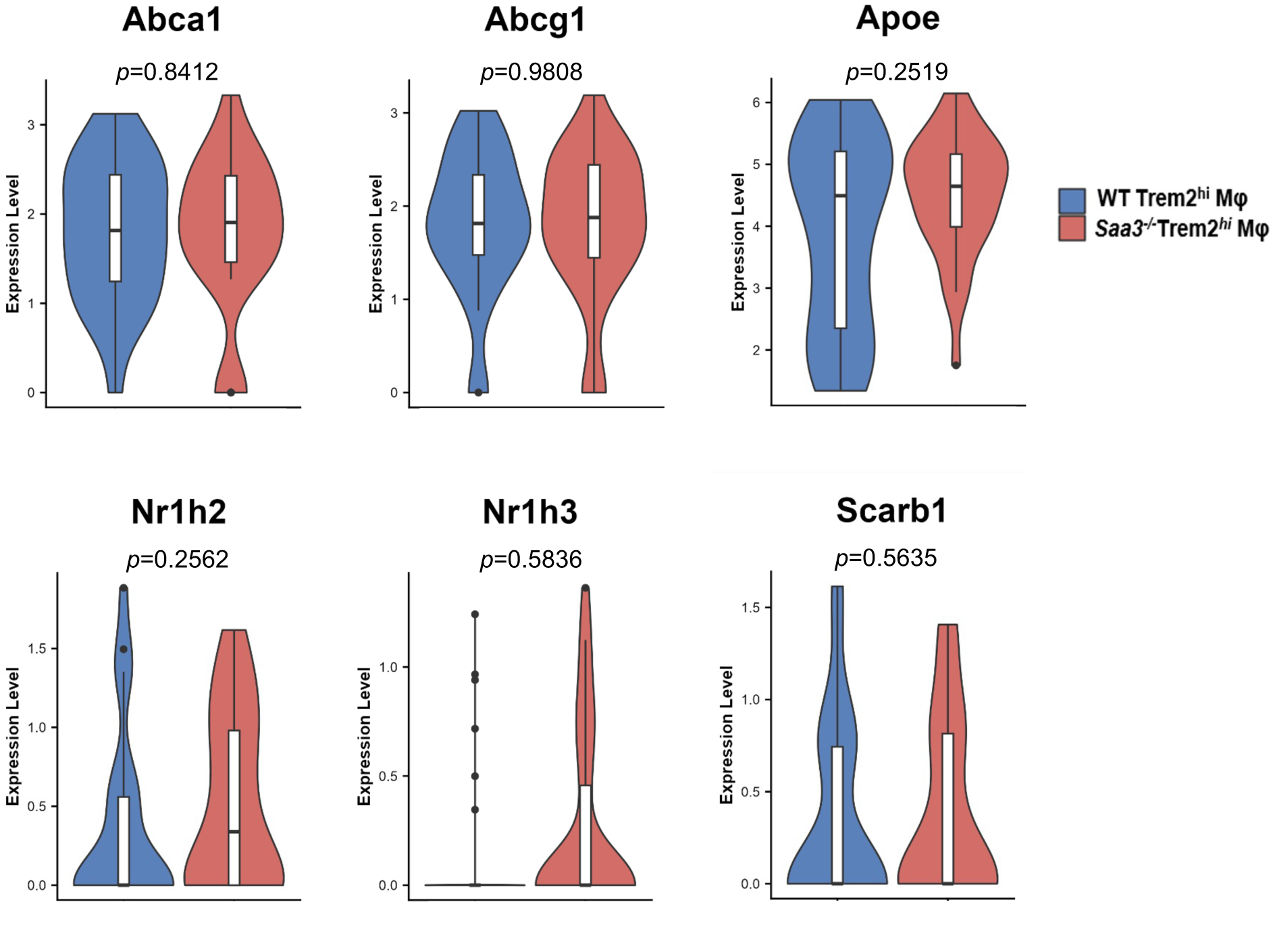
Comparison of differences in the expression level of cholesterol efflux and lipid metabolism genes from Trem2^hi^ macrophages between WT and *Saa3^−/−^* mice. Differential gene expression in aortic Trem2^hi^ macrophages (Mφ) from WT and *Saa3^-/-^* mice. Violin plots illustrate the distribution of expression levels for genes involved in cholesterol efflux and lipid metabolism in Trem2^hi^ macrophages. Blue represents WT macrophages, while red indicates *Saa3^−/−^* macrophages.

